# Chromosome-level reference genome for the medically important Arabian horned viper (*Cerastes gasperettii*)

**DOI:** 10.1101/2024.07.29.605543

**Authors:** Gabriel Mochales-Riaño, Samuel R. Hirst, Adrián Talavera, Bernat Burriel-Carranza, Viviana Pagone, Maria Estarellas, Theo Busschau, Stéphane Boissinot, Michael P. Hogan, Jordi Tena-Garcés, Davinia Pla, Juan J. Calvete, Johannes Els, Mark J. Margres, Salvador Carranza

## Abstract

Venoms have traditionally been studied from a proteomic and/or transcriptomic perspective, often overlooking the true genetic complexity underlying venom production. The recent surge in genome-based venom research (sometimes called “venomics”) has proven to be instrumental in deepening our molecular understanding of venom evolution, particularly through the identification and mapping of toxin-coding loci across the broader chromosomal architecture. Although venomous snakes are a model system in venom research, the number of high-quality reference genomes in the group remains limited. In this study, we present a chromosome-resolution reference genome for the Arabian horned viper (*Cerastes gasperettii*), a venomous snake native to the Arabian Peninsula. Our highly-contiguous genome allowed us to explore macrochromosomal rearrangements within the Viperidae family, as well as across squamates. We identified the main highly-expressed toxin genes compousing the venom’s core, in line with our proteomic results. We also compared microsyntenic changes in the main toxin gene clusters with those of other venomous snake species, highlighting the pivotal role of gene duplication and loss in the emergence and diversification of Snake Venom Metalloproteinases (SVMPs) and Snake Venom Serine Proteases (SVSPs) for *Cerastes gasperettii*. Using Illumina short-read sequencing data, we reconstructed the demographic history and genome-wide diversity of the species, revealing how historical aridity likely drove population expansions. Finally, this study highlights the importance of using long-read sequencing as well as chromosome-level reference genomes to disentangle the origin and diversification of toxin gene families in venomous species.

## Background

The rise of genomics in non-model organisms has led to an increase in the number of high-quality reference genomes available in recent years (Dussex et al., 2021; Hogan et al., 2024; Margres et al., 2021a; Pardos-Blas et al., 2021; Schield et al., 2019; Suryamohan et al., 2020). Such advances in sequencing technologies have catalyzed the study of several complex traits from a genomic perspective, such as coloration, domestication, or venom, among others (Drukewitz & Von Reumont, 2019; Frantz et al., 2020; Margres et al., 2021a; Orteu & Jiggins, 2020; San-Jose & Roulin, 2017). Among these, venom genomic research has been particularly important in enhancing our understanding of the origin, evolution and dynamics of this medically relevant trait (Casewell et al., 2013; Dowell et al., 2016; Giorgianni et al., 2020; Werren et al., 2010). Venom is a potentially lethal cocktail rich in proteins and peptides (from now on referred to as “toxins”) which are actively secreted by specialized venom glands (Casewell et al., 2013; Fry et al., 2009). Toxins can have different effects depending on their type, interactions, and the organism in which they are introduced, with convergent outcomes in different taxa (Fry et al., 2009; Zancolli et al., 2022). Historically, venom research has primarily been conducted using proteomic (and transcriptomic) approaches (see Drukewitz & Von Reumont (2019) and references therein). The identification of venom toxins and the characterization of their evolution using reference genomes is a recent and novel field at all taxonomic levels (Vonk et al., 2013). Previous works have shown that changes in gene regulation can result in the activation and deactivation of venom-coding genes at all taxonomic levels and within the same individual (Avella et al., 2022; Hogan et al., 2024; Margres, Rautsaw, et al., 2021; Zancolli et al., 2022), suggesting that exclusively studying the expression of venom toxins (i.e., proteomics or transcriptomics) is insufficient to disentangle the complete number and biochemical nature of the toxins an individual can potentially transcribe (Drukewitz & Von Reumont, 2019). Ultimately, the study of venom genomics may yield insights into antivenom or drug discovery, as it can identify unexpressed toxin-coding genes that target specific physiological pathways, potentially leading to new therapies for human illnesses including but not limited to cancer (Casewell et al., 2013; King, 2011; L. Li et al., 2018).

Venom has evolved independently in multiple groups including cnidarians, molluscs, arthropods, squamates and even mammals (Casewell et al., 2013; Fry et al., 2009). However, venomous snakes are one of the most life-threatening animal groups to humans (Williams et al., 2019) and, therefore, the fundamental model system in venom research. Venomous snakes are a diverse group with more than 600 species (Uetz, 2021), where venom has evolved with the objective of immobilizing and digesting their prey (Fry & Wüster, 2004). From those, more than 370 species have been classified as of medical importance by the World Health Organization (WHO) due to their potential severe effects on humans. In fact, snakebite is considered a neglected tropical disease, with annual mortality exceeding 100,000 victims worldwide (Gutiérrez et al., 2017; Williams et al., 2019). Within venomous snakes, the most medically important families are Elapidae, Viperidae and Atractaspididae (Tasoulis & Isbister, 2017), although within Colubridae (*sensu lato*) there are certain medically important venomous species as well (Weinstein et al., 2013). Envenomation by certain members of these families can result in a range of pathologies, spanning neurotoxic, hemotoxic, and/or cytotoxic effects (among others) depending on the number and composition of toxins. Neurotoxic venoms primarily target the central nervous system and are mainly composed of small proteins including three-finger toxins (3FTs), snake venom phospholipases A_2_ group I (SV-G^I^-PLA_2_) or dendrotoxins, and are usually associated with elapid snakes (Ferraz et al., 2019). Conversely, hemotoxic and cytotoxic venoms generally are comprised of large enzymatic proteins and protein complexes, including snake venom metalloproteases (SVMP), serine proteases (SP) or snake venom phospholipases A_2_ group II (SV-G^II^-PLA_2_), and are typically associated with viperid snakes (Fry, 2015; Fry et al., 2008; Tasoulis & Isbister, 2017). While these historical classifications have proven to be somewhat useful for treating envenomations medically, recent studies have revealed that the presence of these toxins are not exclusive to specific snake families (Osipov & Utkin, 2023).

Vipers (family Viperidae) are a monophyletic lineage of venomous snakes found across Eurasia, Africa and America (Vitt & Caldwell, 2014), receiving extensive research attention primarily due to their medical relevance (Arnold et al., 2009; Casewell et al., 2009; Pook et al., 2009; Šmíd & Tolley, 2019; Wüster et al., 2008). The majority of venom studies in this group have primarily been conducted using a proteomic approach, with early venom work being highly motivated by the medical field, with a limited number of studies employing genomic approaches (but see Almeida et al., (2021); Margres et al., (2021a); Myers et al., (2022); Schield et al., (2019); Hirst et al., (in review); Hogan et al., (2024)). Sequencing efforts to obtain high-quality reference genomes have mainly focused on pitvipers (Crotalinae subfamily, 11 reference genomes, NCBI last accessed 13 March 2024), especially within the *Crotalus* (*n*=6) genus, and have focused on the study of venom evolution (Gilbert et al., 2014; Hogan et al., 2021; Margres et al., 2021a; Schield et al., 2019; Westeen et al., 2023). Other viperids have also been sequenced (although in lower numbers) from both Azemiopinae and Viperinae subfamilies (one and four, respectively) (Myers et al., 2022; Saethang et al., 2022; Talavera et al., in review). Currently, a total of 16 species within the Viperidae family posses an available reference genome at NCBI, corresponding to 3.6% of the total 387 species (Uetz, 2021). Vipers display extensive variation in venom composition between and within genera (Ali et al., 2015; Mackessy, 2010) and even intraspecifically (Jan et al., 2002; Zancolli et al., 2019). Such differences are most likely due to the high diversity of venom genes and their different effects on prey but also, at least on some occasions, the result of introgression with related species (Jan et al., 2002; Margres et al., 2021b; Smith et al., 2023). This provides an extraordinary opportunity to study trait evolution both at the inter- and intraspecific levels.

Native to the Arabian Peninsula, the Arabian horned viper (*Cerastes gasperettii*, family Viperidae) is a venomous snake currently recognized within the highest medical importance category (WHO; accessed July, 2024). Extending from the Sinai Peninsula to southwestern Iran in the north and reaching as far as Yemen and Oman in the south, its distribution is widespread (Fig. S1). Found mainly in sandy habitats, this arid-adapted ground-dwelling snake with generalist requirements (Carranza et al., 2021; Mochales-Riaño et al., 2024; Russell & Campbell, 2015) is one of the most common venomous snakes found in Arabia and is responsible for occasional snakebite envenomations (Al-Sadoon & Paray, 2016; Amr et al., 2020; Schneemann et al., 2004).

In this study, we present a high-quality chromosome-level reference genome assembly for the Arabian horned viper (*Cerastes gasperettii*, NCBI: txid110202), being one of the first within the Viperinae subfamily. Our highly-contiguous genome showcases a high level of similarity at the chromosome level within the Viperidae family with some minor rearrangements with elapids. Moreover, employing several -omics techniques, we characterized the main toxins found in its venom and the location of those toxins in the genome, comparing their evolutionary history and gene copy number variation with other venomous species. We deciphered numerous genomic attributes of this species including its genetic diversity and failed to find evidence of inbreeding. Finally, we reconstructed the demographic history for the species, revealing how historical increases in aridity likely drove population expansions. Overall, the genomic resources generated in this study provide an essential reference resource for forthcoming studies on venom evolution.

## Methods

### Sampling

Three adult specimens (two females and one male) of *Cerastes gasperettii gasperettii* were used for this study (Table S1). Blood was extracted only from a single female individual (the heterogametic sex, sample CG1) to obtain High-molecular-weight (HMW) genomic DNA (gDNA) and stored in ethanol and EDTA. For each of the three individuals, we extracted twelve different tissues, including the venom gland, which was stored in RNAlater until RNA extraction (Table S1 and Fig. S2). We only extracted the left venom gland per individual, as previous research within the same family has shown that both venom glands provide indistinguishable results (Rokyta et al., 2017).

### DNA extraction, library preparation and sequencing

We extracted gDNA from the blood of a female individual (CG1 in Table S1) using the MagAttract HMW Kit (Qiagen) following manufacturer’s protocols. Then, we sequenced a total of two 8M SMRT HiFi cells, aiming for a ∼30x of coverage, at the University of Leiden. Hi-C libraries were prepared using the Omni-C kit (Dovetail Genomics), following the manufacturer’s protocol and using blood stored in EDTA, at the National Center for Genomic Analyses (CNAG), in Barcelona, Spain. The library was paired-end sequenced on a NovaSeq 6000 (2 × 150 bp) following the manufacturer’s protocol for dual indexing and aiming for a coverage of ∼60x. Finally, we sequenced short-read whole-genome data of the same individual using a NEB Ultra II FS DNA kit; the library was paired-end sequenced on a NovaSeq 6000 (2 × 150 bp) at the Core sequencing platform from the New York University of Abu Dhabi, aiming for ∼70x depth of coverage.

### RNA extraction, library preparation and sequencing

We extracted RNA from the same three individuals described above (Table S1 and Fig. S2). RNA was isolated using the HighPurity^TM^ Total RNA Extraction Kit (Canvax, Valladolid, Spain). We selected a total of 35 samples (including venom glands, tongue, liver and pancreas, among others; Table S2). RNA libraries were prepared with the VAHTS Universal V8 RNA-seq Library Prep Kit and were sequenced on a NovaSeq 6000 (2 × 150 bp) aiming for an average of 40M read pairs per sample (Table S2). Moreover, we sequenced one 8M SMRT HiFi cell containing two Iso-seq HiFi libraries at University of Leiden: one containing only the venom gland, and the second library being a pool of eight high-quality tissues (brain, kidney, liver, gallbladder, spleen, tongue, pancreas and ovary).

### Genome assembly and scaffolding

Quality control on HiFi and Illumina reads was assessed using FastQC (Andrews, 2010) and adapters were removed with cutadapt (Martin, 2011). To make an initial exploration of the genome, using the raw HiFi reads, we generated a k-mer profile with Meryl (Rhie, Walenz, et al., 2020) and visualized it with GenomeScope2 (Ranallo-Benavidez et al., 2020). Then, we assembled the genome following the VGP assembly pipeline v2.0 (Rhie, McCarthy, et al., 2020). PacBio HiFi reads were assembled into contigs using the software Hifiasm (Cheng et al., 2021), producing primary and alternate assemblies. We used *purge_dups* (Guan et al., 2020) to remove haplotypic duplicates from the primary assembly and added them to the alternate assembly. Then, we scaffolded the primary assembly using the Hi-C data with SALSA2 (Ghurye et al., 2019). Manual curation was performed with <u>Pretext</u>. We used the ∼78x Illumina data to polish the assembly with one round of Pilon (Walker et al., 2014). The mitochondrial genome was obtained with GetOrganelle (Jin et al., 2020), using the mitochondrial genome of several *Echis* species (*E. coloratus, E. carinatus* and *E. omanensis*) to seed the assembly (NCBI accession numbers: <u>SRX18902082, SRX18902083</u>, <u>SRX18902084</u>, respectively).

### Genome assembly quality evaluation

Quality assessment and general metrics for the final assembly were estimated with both QUAST v.5.1.0 (Gurevich et al., 2013) and gfastats (Formenti et al., 2022). Possible contaminations were evaluated with BlobToolKit (Challis et al., 2020) using the NCBI taxdump database. We also used MitoFinder (Allio et al., 2020; D. Li et al., 2016) to confirm that the mitochondrial genome was absent in the assembled nuclear reference genome. Completeness of the genome assembly was assessed with BUSCO v5.3.0. against the sauropsida_odb10 database (*n*=7,480).

### Genome annotation

First, we identified repetitive elements using RepeatModeler v.2.0.3 (Flynn et al., 2020) for *de novo* predictions of repeat families. To annotate genome-wide complex repeats, we used RepeatMasker v.4.1.3 (Tempel, 2012) with default settings to identify known Tetrapoda repeats present in the curated Repbase database (Bao et al., 2015). Then, we ran three iterative rounds of RepeatMasker to annotate the known and unknown elements identified by RepeatModeler and soft-masked the genome for simple repeats. We used GeMoMa v.1.9 (Keilwagen et al., 2019) to annotate protein-coding genes, combining both the RNA-seq data generated in this study as described above as well as annotations from seven other squamate genomes already published: *Anolis carolinensis* from Alföldi et al., (2011), *Crotalus adamanteus* from Hogan et al., (2024), *Crotalus tigris* from Margres et al., (2021a), *Ophiophagus hannah* from Vonk et al., (2013), *Naja naja* from Suryamohan et al., (2020), *Crotalus ruber* from Hirst et al., (in review) and *Crotalus viridis* from Schield et al., (2019). We previously quality checked and removed the adapters of the RNA-seq data as well as mapped the transcriptomic data to our new reference genome with Hisat2 (Kim et al., 2019). Additionally, we also removed the adapters for the Iso-seq data and mapped the long-read transcriptomic data to our new reference genome with pbmm2, collapsing mapped reads into unique isoforms with isoseq3 and annotated with GeneMarkS-T (S. Tang et al., 2015). We combined both annotations (GeMoMa and GeneMarkS-T) with TSEBRA (Gabriel et al., 2021). We blast our predicted proteins to a Uniprot protein database for a total of ten species (*C*. *gasperettii*, *C. vipera*, *C. cerastes*, *Anolis carolinensis*, *Crotalus viridis*, *Crotalus tigris*, *Crotalus ruber*, *Crotalus adamanteus*, *Ophiophagus hannah* and *Naja naja*). Simultaneously, we ran Interproscan (Jones et al., 2014) on our predicted proteins. Then, we combined both functional annotations with AGAT (Dainat et al., 2023). Finally, as venom-gene families are known to occur in large tandem arrays and the number of paralogs can be underestimated in particular gene families (Schield et al., 2019), we performed additional annotation steps for venom genes. Following Margres et al., (2021a), we used a combination of empirical annotation in FGENESH+ (Solovyev et al., 2006), as well as manual annotation using RNA-seq and Iso-seq alignments; the former identified all genes regardless of expression, whereas the latter was used to explicitly identify expressed toxins.

### Macrosynteny analyses

Whole-genome synteny was explored between our new chromosome-level reference genome for the Arabian horned viper together with the Eastern diamondback rattlesnake (*Crotalus adamanteus*) (Hogan et al., 2024), the Indian cobra (*Naja naja*) (Suryamohan et al., 2020) and the Brown anole (*Anolis sagrei*) (Geneva et al., 2022) using MCscan (H. Tang et al., 2008). Protein sequences from each of the three venomous snakes were extracted using AGAT (v1.2.1) (Dainat et al., 2023) and were pairwise aligned with LAST (Kiełbasa et al., 2011), implemented in the JCVI python module (Tang et al., 2017). A first alignment was used between the three species to identify chromosomes assembled in the reverse complement, which were corrected using SAMtools faidx (v1.18.1) (Danecek et al., 2021). Gene annotations for the new reference (with the corresponding reversed chromosomes) were annotated using GeMoMa v.1.9 (Keilwagen et al., 2019), and MCscan was rerun.

### Transcriptomics

After adapter trimming and quality control, we mapped our RNA-seq reads to the reference genome of *Cerastes gasperettii* using Hisat2 (Kim et al., 2019). Gene counts per gene across all samples were calculated with StringTie (Pertea et al., 2015). Initial exploration of our transcriptomic data revealed a clear batch effect for one of the three samples (Fig. S4), due to the low mapping of that sample to our reference genome. Therefore, we decided to remove individual CG1 from future RNA-seq analyses. Moreover, to avoid pseudoreplication, we also removed the accessory gland from individual CG009 due to its high similarity with the venom gland, suggesting that the venom gland rather than the accessory gland was sampled (Fig. S4). Differential expression analyses were carried out with the DESeq2 package (Love et al., 2014) from R (R Core Team, 2021), using the DESeq2 median of ratios normalization. Finally, we identified the highly expressed genes found in the venom gland as well as the toxins uniquely expressed in the venom gland (following Suryamohan et al., (2020)) which were defined as (1) genes expressed in the venom gland (TPM > 500), (2) Differential Upregulated Genes (DUGs) with Fold Change (FC) > 2 comparing venom glands to all other tissues and (3) unique to venom glands (TPM < 500 in all other tissues).

### Proteomics

A bottom-up mass spectrometry strategy (Calvete et al., 2021) was used to characterize the venom arsenal of *Cerastes gasperettii*. Briefly, the venom proteome (pool from individuals CN6134 and CN6135; Table S1) was submitted to reverse-phase High-performance liquid chromatography (HPLC) decomplexation followed by SDS-PAGE analysis in 12% polyacrylamide gels run under non-reducing and reducing conditions. Protein bands were excised from Coomassie Brilliant Blue-stained gels and subjected to automated in-gel reduction and alkylation on a Genomics Solution ProGest™ Protein Digestion Workstation. Tryptic digests were submitted to MS/MS analysis on a nano-Acquity UltraPerformance LC^®^ (UPLC^®^) equipped with a BEH130 C_18_ (100µm x 100mm, 1.7 µm particle size) column in-line with a Waters SYNAPT G2 High Definition mass spectrometer. Doubly and triply charged ions were selected for CID-MS/MS. Fragmentation spectra were matched against a customized database including the bony vertebrates taxonomy dataset of the NCBI non-redundant database (release 258 of October 15, 2023) plus the species-specific venom gland transcriptomic and genomic protein sequences gathered in this work. Search parameters were as follows: enzyme trypsin (two-missed cleavage allowed); MS/MS mass tolerance for monoisotopic ions: ± 0.6 Da; carbamidomethyl cysteine and oxidation of methionine were selected as fixed and variable modifications, respectively. Assignments with significance protein score threshold of *p* < 0.05 (Mascot Score > 43) were taken into consideration, and all associated peptide ion hits were manually validated. Unmatched MS/MS spectra were *de novo* sequenced and manually matched to homologous snake venom proteins available in the NCBI non-redundant protein sequences database using the default parameters of the BLASTP program (https://blast.ncbi.nlm.nih.gov/Blast.cgi).

### Microsynteny

To explore toxin genomic organization across (sub)families, we used blast, incorporating both toxin and non-toxin paralogs to identify the genomic location of SVMPs, SVSPs and PLA_2_ toxin families, across the genome of *Cerastes gasperettii*, *Crotalus adamanteus*, *N. naja* and *A. feae*. We excluded *A. feae* for SVSPs and SVMPs microsynteny analyses as those families were not assembled onto a single contig in the *A. feae* genome. Then, we aligned those regions using Mafft (Katoh & Standley, 2013). Each species was annotated within the MSA using its own annotation as a reference in Geneious Prime 2023.0.4. Results were plotted using the gggenomes package (https://github.com/thackl/gggenomes) from R (R Core Team, 2021).

### Toxin phylogenies

We used phylogenetic inference to study the evolutionary history for the main groups of toxins (i.e., SVMPs and SVSPs, which were the most abundant in the proteome of *Cerastes gasperettii*, as well as PLA_2_ as this family has been widely studied within the Viperidae family (Dowell et al., 2016; Myers et al., 2022). For the three main toxin families, we selected available toxin genes as well as non-toxin paralogous genes from venomous species; we also included other non-toxin paralogous genes from non-toxic species (for details about this see Supplementary Information). When needed, we translated CDS to protein sequence, and then protein sequences were aligned with Mafft (Katoh & Standley, 2013). Following Giorgianni et al., (2020), we built a phylogeny for each of the toxin groups using Phyml v3.3 (Guindon et al., 2010), implementing the Dayhoff substitution model and validating our inferred tree with aBayes support.

### Demographic history

We inferred the demographic history of *Cerastes gasperettii* by implementing the Pairwise Sequential Markovian Coalescent (PSMC) software (H. Li & Durbin, 2011) on the short-read whole-genome data. Heterozygous positions were obtained from bam files with Samtools v1.9 (H. Li et al., 2009), and data were filtered for low mapping (<30) and base quality (<30). Minimum and maximum depths were set at a third (27x) and twice (156x) the average coverage. Only autosomal chromosomes were considered. We used the squamate mutation rate of 2.4×10^-9^ substitutions/site/generation and a generation time of 3 years, following Green et al., (2014) and Schield et al., (2022), respectively. A total of ten bootstraps were calculated, plotting the final results with the psmc_plot.pl function from PSMC (https://github.com/lh3/psmc).

### Genomic diversity

We downloaded Illumina data for *Bothrops jararaca* (SRR13839751 from Almeida et al., (2021)), *Crotalus viridis* (SRR19221440 from Schield et al., (2019)), *Naja kaouthia* (SRR 8224383; Thongchum et al., (2019)), *Naja naja* (SRR10428156; Suryamohan et al., (2020)) and *Sistrurus tergeminus* (SRR12802282; Bylsma et al., (2022)). Then, we filtered for quality (Phred score of 30) and removed adapters with fastp (Chen et al., 2018). Trimming of poly-G/X tails and correction in overlapped regions were specified. All other parameters were set as default. Filtered sequences were visually explored with FastQC (Andrews, 2010) to ensure data quality and absence of adapters. Filtered reads were mapped against the new reference genome of *Cerastes gasperettii* using the bwa mem algorithm (H. Li, 2013). Mapped reads were sorted with Samtools v1.9 (H. Li et al., 2009) and duplicated reads were marked and removed with PicardTools (Broad Institute, 2021). Reads with mapping quality lower than 30 were discarded. SNP calling was carried out with HaplotypeCaller from GATK (McKenna et al., 2010), with BP_resolution and split by chromosome. For each chromosome, individual genotypes were joined using CombineGVCFs with convert-to-base-pair-resolution, and the GenotypeGVCFs tool was then applied to include non-variant sites. Finally, for each individual, the whole dataset split by chromosome was concatenated with bcftools concat (Danecek et al., 2021), keeping only the autosomes. Then, for each sample, we used the raw dataset to calculate average genome heterozygosity. We generated non-overlapping sliding windows of 100 Kbp for the newly assembled *Cerastes gasperettii* reference genome and included only sites (both variant and invariant) with site quality higher than 30 (QUAL field in a VCF file from GATK). Only windows containing more than 60,000 unfiltered sites were considered. Visualization was carried out with ggplot2 (Wickham, 2016) in R (R Core Team, 2021).

## Results and Discussion

### Genome assembly and annotation

We generated a high-quality chromosome-level assembly for the Arabian horned viper (*Cerastes gasperettii*) by combining PacBio HiFi (∼40x), Hi-C (∼60x) and Illumina data (∼78x) (Fig. 1 and Fig. S3). First, we *de novo* assembled the HiFi reads into 1,018 contigs (N50=45.7 Mbp; longest contig of 149.99 Mbp). Then, using the proximity ligation data (i.e., Hi-C), we scaffolded the genome into 319 scaffolds (N50=111.38 Mbp; largest scaffold 345.38 Mbp). After manual curation, we enhanced the scaffolding parameters of our genome (N50=214.14 Mbp; largest scaffold 361.99 Mbp), containing 99.44% of the genome present in 19 scaffolds or pseudochromosomes (7 macro-, 10 micro-, Z and W sex chromosomes; Table 1 and Fig. 1B). The total genome length was 1.63 Gb, similar to other venomous snakes (Margres et al., 2021a; Schield et al., 2019; Suryamohan et al., 2020; Vonk et al., 2013; Table 1), with a contig N50 of 45.6 Mbp, ∼3.3 times more contiguous than the *N. naja* genome (Suryamohan et al., 2020), ∼228 times more contiguous than the *Anolis sagrei* genome (Geneva et al., 2022), but 0.67 times less contiguous than the recently published *Crotalus adamanteus* genome (Hogan et al., 2024), making it one of the most contiguous chromosomal squamate genomes assembled to date (Table 1). We assessed the completeness of the assembly using BUSCO (Simão et al., 2015) with the sauropsida gene set (*n*=7,480). Upon evaluation, we successfully identified 92.8% of the genes (91.4% single-copy, 1.4% duplicated), while the remaining genes were fragmented (1%) or missing (6.2%; Fig. 1). For the *de novo* assembly, GC content and repeat content were 37.87% and 43.63%, respectively. The repetitive landscape was dominated by retroelements (30.25%), with a majority of LINEs (21.25%) (Table S3). Finally, we annotated 27,158 different protein-coding genes within our assembly, with a total of 194 putative toxins or toxin-paralogs genes. Toxin genes usually found in venomous snakes (see proteome results below) were mainly found on macrochromosomes, although major toxin groups were found on microchromosomes (SVMPs, SVSPs and PLA_2_; Fig. 1). Finally, we also found a battery of 3FTxs and myotoxin-like genes, but they were not represented in our RNA-seq dataset (see below).

**Fig. 1:**
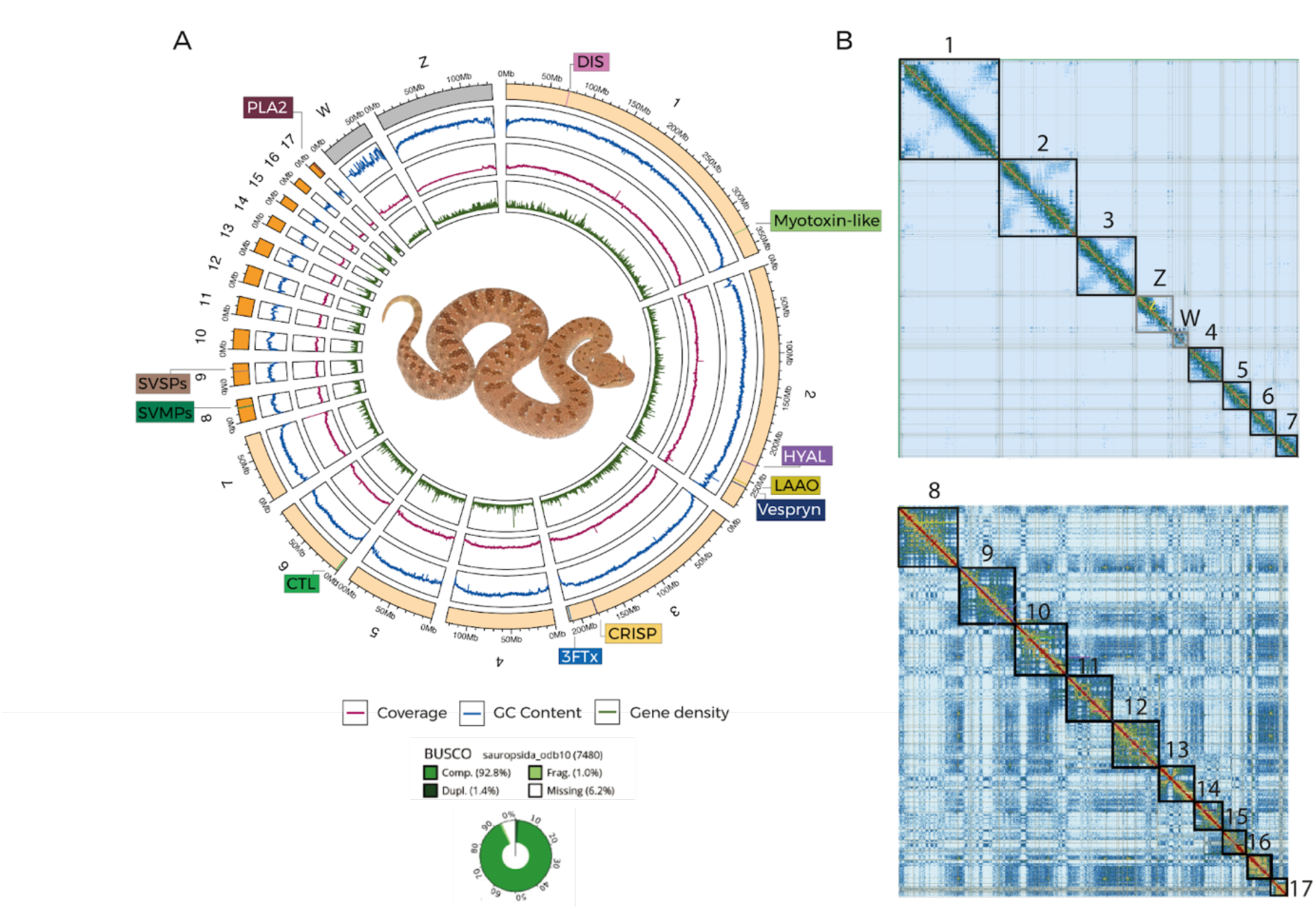
A) Reference genome for *Cerastes gasperettii*, including BUSCO score, GC content, coverage level as well as the main toxins found within the genome. Macrochromosomes are shown in light orange whilst microchromosomes are shown in bright orange. Sex chromosomes are shown in gray. Abreviations are as follows: DIS, Disintegrins; HYAL, Hyaluronidases; LAAO, L-Amino acid oxidase; CRISP, Cysteine-rich secreted proteins; 3FTx, Three-finger toxins; CTL, C-type lectins; SVMPs, Snake venom metalloproteinases; SVSPs, Snake venom serine proteinases; PLA2, Phospholipases. B) Linkage map for the macrochromosomes (above), including the sex chromosomes (Z and W), and microchromosomes (below).

**Table 1:** Comparison of our new reference genome for *Cerastes gasperettii* with other high-quality squamate genomes. Best value per category is shown in bold.

|  | *Cerastes gasperettii* | *Crotalus adamanteus* | *Naja naja* | *Anolis sagrei* |
| --- | --- | --- | --- | --- |
| Genome size | 1.63 Gbp | 1.69 Gbp | 1.79 Gbp | 1.92 Gbp |
| Number of scaffolds | 221 | **27** | 1,897 | 3,738 |
| Scaffold N50 | 214.14 Mbp | 208.9 Mbp | 223.35 Mbp | **253.58 Mbp** |
| Scaffold L50 | **3** | **3** | **3** | 4 |
| Contig N50 | 45.6 Mbp | **67.5 Mbp** | 13.06 Mbp | 0.2 Mbp |

### Macrosynteny

Whole-genome synteny comparisons showed a great level of similarity between *Cerastes gasperettii* and *Crotalus adamanteus*, with large syntenic blocks both within macro- and microchromosomes (Fig. 2). Some chromosomal rearrangements were observed between viperids and elapids, as previously discussed by Suryamohan et al., (2020), with a fission of chromosome four in *N. naja* to form chromosomes five and seven in vipers, and a fusion of chromosomes five and six in *N. naja* to form chromosome four in vipers. Interestingly, several chromosomal rearrangements between lizards and snakes have occurred, as we found several fission events in the *A. sagrei* genome, including one fission from chromosome two to originate the current Z chromosome in snakes (Fig. 2). The last four scaffolds (14, 15, 16 and 17) from *Anolis sagrei* were removed, as no orthologous groups were found. Macrosyntenic differences between lizards and snakes could be related to the innovations in different areas such as locomotion, feeding and sensory processing that snakes experienced during their origin more than 150 Mya (Title et al., 2024) as well as explain the high level of chromosomal similarity within snakes.

**Fig. 2:**
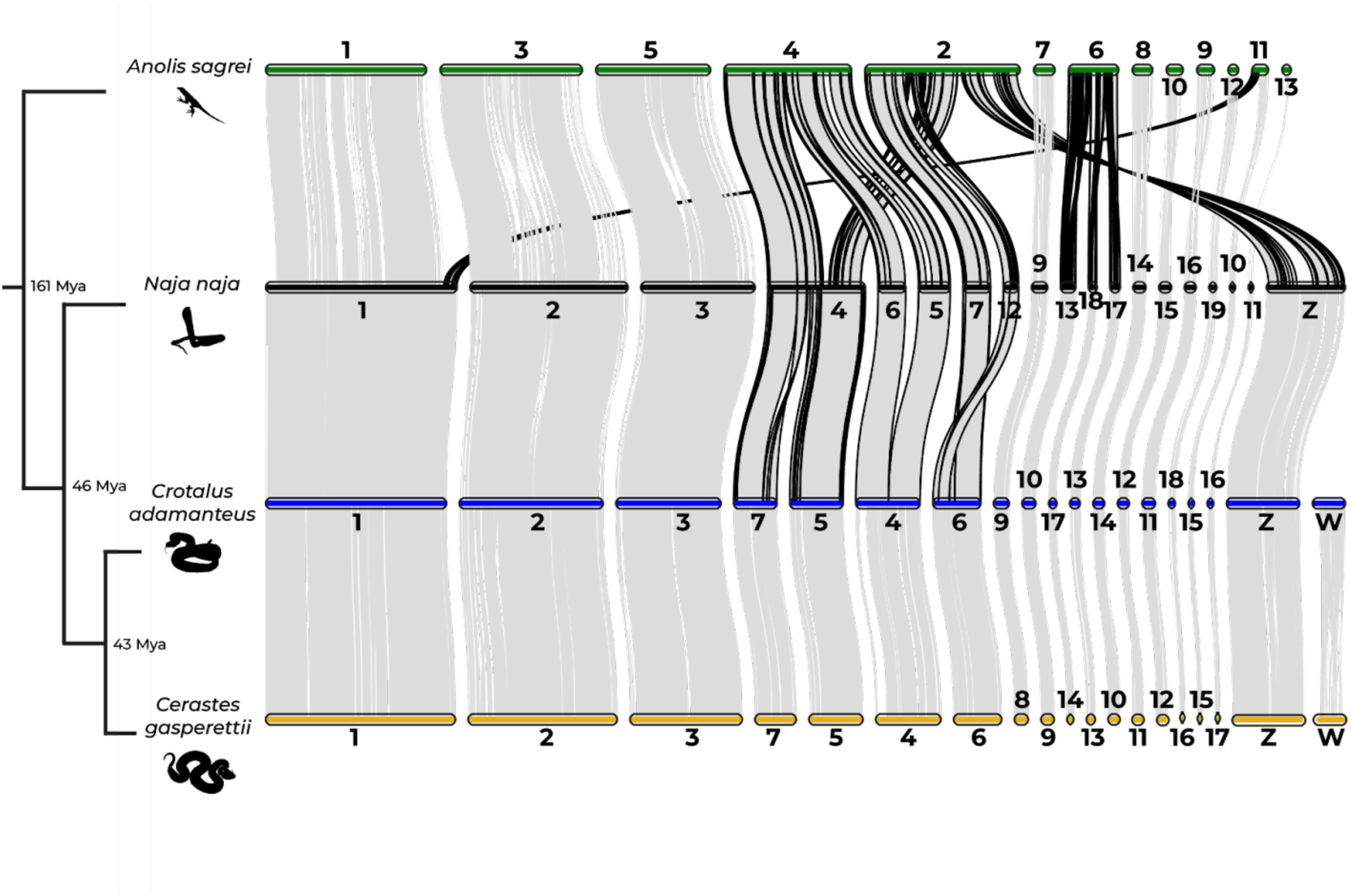
Macrosynteny analyses for one Elapidae (*Naja naja*), one Crotalinae (*Crotalus adamanteus*) and one Viperinae (*Cerastes gasperettii*) species, with *Anolis sagrei* as the outgroup. The four smallest scaffolds (14, 15, 16 and 17) of *Anolis sagrei* were removed, as no orthologous groups were found with other species. Borders of regions showing evidence for chromosomal rearrangements are shown in black. Estimates for branch times obtained from TimeTree.org based on divergence times between Iguania and Serpentes, Elapidae and Viperidae and Crotalinae and Viperinae, respectively.

### Transcriptomics

Our analyses of multi-tissue transcriptomic data (23 samples from two individuals covering 13 different tissues) reported a total of 23,178 expressed genes (TPM > 1). Heatmap analyses with the most 2,000 variable genes reported unique upregulated genes for each of the different analyzed tissues (Fig. S5). The venom gland transcriptome contained a total of 7,237 genes expressed (TPM > 500), including a total of 65 putative toxin genes. Differential gene expression analyses revealed a total of 161 genes (33 putative toxin genes) that were differentially upregulated (FC > 2 and 1% FDR) in venom glands compared to other tissues (Fig. 3A). Finally, a total of 10 toxin genes (*CRISP2*, *SVMP9*, *SVMP10*, *SVSP8*, *SVSP7*, *SVSP5*, *CTL14*, *CTL15*, *SVSP4* and *SVMP13*) were uniquely expressed in the venom gland, encoding for the minimal core venom effector (Fig. 3A) (Suryamohan et al., 2020), and in line with the main toxins found within the proteome (Fig. 3B). These genes, together with other SVMPs, SVSPs, Disintegrins (DISI) and C-type lectins (CTL), were highly expressed in the venom gland and form the core toxic effector components of the venom. Targeting the core venom toxins together with other well-known modulators of venom may help manufacture of synthetic antivenom treatments as well as improve neutralization tests of current antivenoms (Suryamohan et al., 2020). However, more transcriptomic data should be incorporated to correct for potential ontogenetic and geographical variation in venom composition in *C. gasperettii* (Avella et al., 2022; Kalita et al., 2018).

**Fig. 3:**
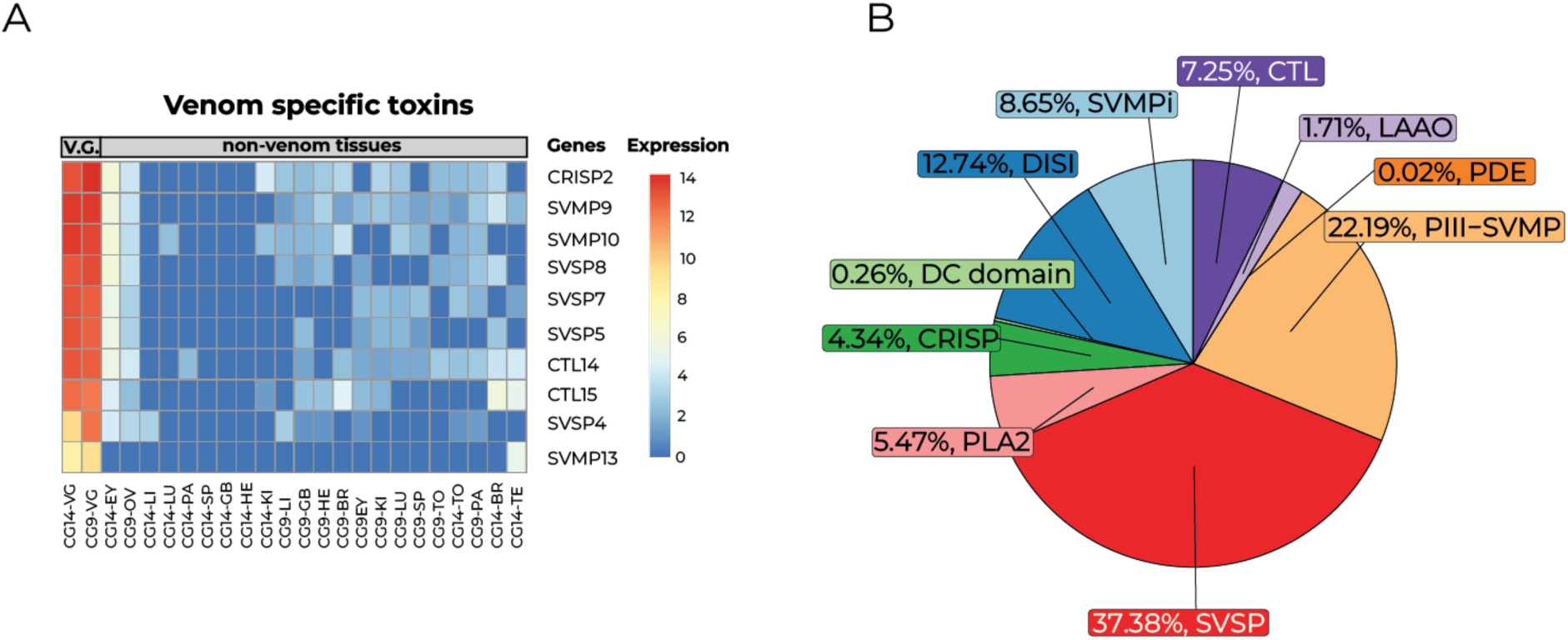
Main toxins found in both the transcriptome and proteome of *Cerastes gasperettii*. A) Genes upregulated and exclusively found in the venom gland for both individuals. Each column represents a different tissue type per sample. Rows show the different genes, and colors correspond to different expression levels. Abbreviations are as follows: VG, Venom Gland; EY, Eye; OV, Ovary; LI, Liver; LU, Lung; PA, Pancreas; SP, Spleen; GB, Gallbladder; HE, Heart; KI, Kidney; LI, Liver; BR, Brain; TO, Tongue; TE, Testis. B) Venom composition for one individual of *Cerastes gasperettii*. The pie chart displays the relative abundances of the toxin families found in the proteome of the *Cerastes gasperettii* venom. Abbreviations are as follows: SVMP, snake venom metalloproteinase; SVSP, snake venom serine proteases; PLA2, phospholipases A2; CRISP, cysteine-rich secretory proteins; DISI, disintegrins; CTL, C-type lectins; LAAO, L-amino-acid oxidases; PDE, phosphodiesterases.

### SVSPs and SVMPs as main toxin proteins

Venom proteomics identified SVSPs and SVMPs as the most abundant toxin families within the venom of *Cerastes gasperettii*, with 37.38% and 22.19% of the venom being composed by peptides from those two families, respectively (Fig. 3B); the dominance of these two toxin families is consistent with previous research on the same genus (Casewell et al., 2014; Fahmi et al., 2012). Other toxin families identified were DISI (12.74%), CTL (7.25%), PLA_2_ (5.47%), Cysteine-Rich Secretory Proteins (CRISP; 4.34%) or L-Amino acid oxidase (LAAO; 1.71%) (Fig. 3B).

### SVMPs

We studied venom evolution within the most abundant toxin groups (i.e., SVMPs and SVSPs, as well as PLA_2_). After a thorough manual curation, we used comparative genomics to evaluate the number and position of those genes in comparison with the Indian cobra (*N. naja*), the Eastern diamondback rattlesnake (*Crotalus adamanteus*), and the Fea’s viper (*A. feae*). We reported a total of 13 fully contiguous tandem repeat SVMPs for *Cerastes gasperettii* (Fig. 4A), next to the non-toxic paralogous gene *ADAM28* and flanked by the *NEFL* and *NEFM* non-toxic genes. Microsyntenic analyses showed gene copy number variation between the studied species (Fig. 4A). Overall, we can see an expansion in the number of SVMPs within the Viperidae family, particurlarly in *Crotalus adamanteus* but also in *Cerastes gasperettii* (Fig. 4A). The amplification of SVMP copy numbers is consistent with our proteomic results, as SVMPs were the second most abundant component of the venom (Fig. 3B). Then, we reconstructed the evolutionary history of this toxin family (Fig. 4B and Fig. S6). Phylogenetic analyses for this toxin group reported a highly supported clade comprising *ADAM28* peptides, the non-toxic paralogous gene. The second clade of orthologous toxin-peptides were found within both elapid and viperid families (including species from Crotalinae and Viperinae subfamilies in viperids; Fig. S6) as well as two SVMPs from *A. feae*. Interestingly, we report a new toxin gene within *Cerastes gasperettii* with a different evolutionary history, as it did not share orthology with any other gene (Fig. 4B). This new gene likely arose from a duplication event of *SVMP13*, within the group of SVMP *MDC1* toxins (Fig. S6). This discovery highlights the importance of using genomics in studying venom evolution, as this putatively toxic gene was not found to be differentially upregulated in the venom gland or recovered in the proteome (Fig. 3). More genomic data will indicate if *SVMP12* is unique for the Viperinae subfamily, the *Cerastes* genus or if it is only found in *Cerastes gasperettii*. All other clades were unique to viperids (and some exclusive only to crotalids), except for a clade composed by SVMPs unique to elapids, as previously discussed in Suryamohan et al., (2020). Interestingly, one of the toxins (*SVMP8*) was not a class P-III SVMP, as it within the MAD-4/5 clade (class P-II SVMP), contrary to the proteomic results where all SVMPs were categorized within the class P-III (Fig. 3B). Although there has been a clear expansion of the SVMP family within the *Crotalus* genus, our results suggest that the origin of that expansion was at the beginning of the Viperidae family, as most of the groups are also present within the Viperinae subfamily.

**Fig. 4:**
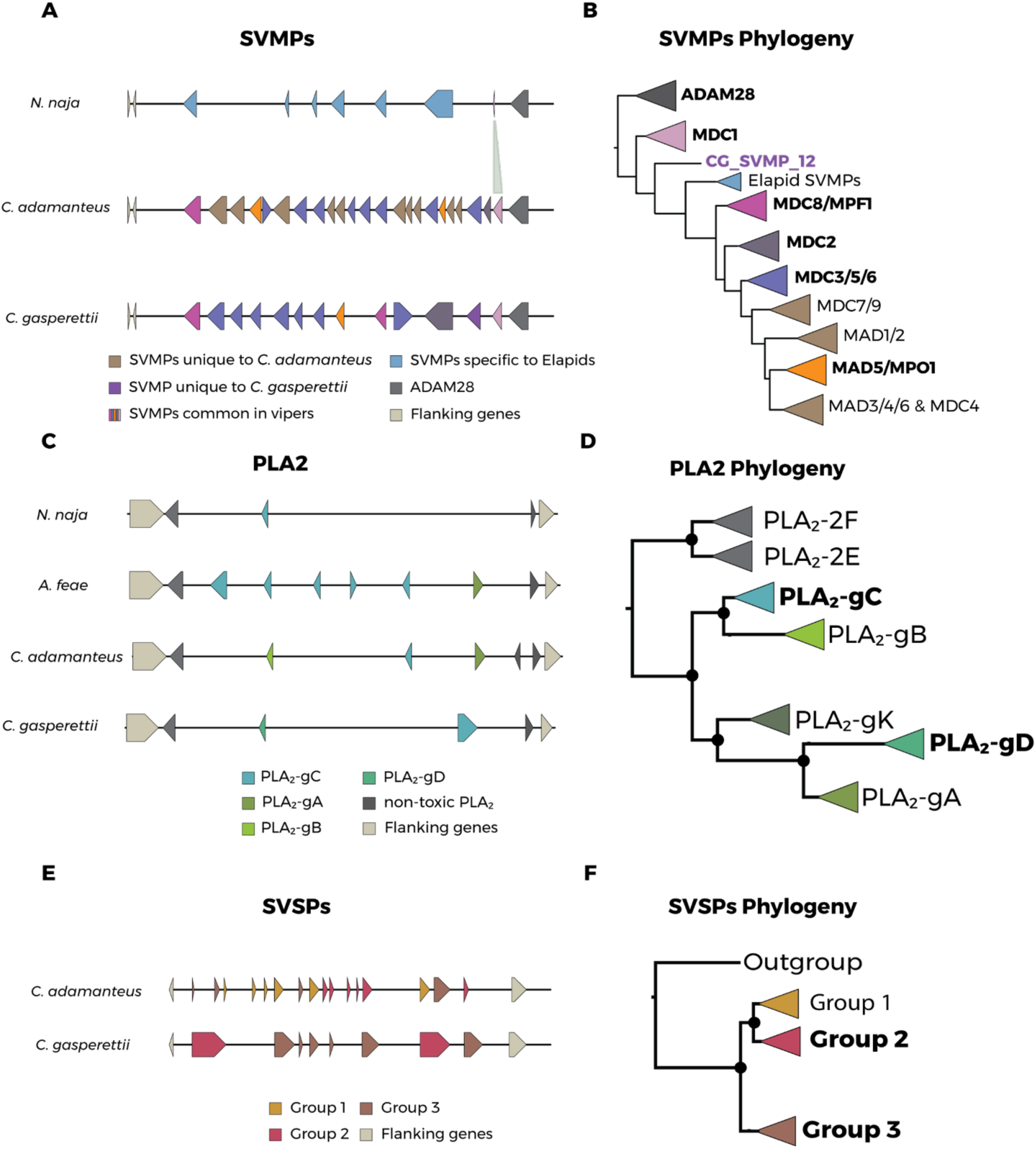
A) Microsynteny for the SVMP toxin family in *Naja naja*, *Crotalus adamanteus* and *Cerastes gasperettii*. Different colors indicate genes unique to *C. gasperettii*, crotalids, true vipers or elapids. ADAM28 (right) as well as flanking genes (left) are also indicated. B) Phylogeny of SVMPs, In bold, groups that contained SVMPs from *Cerastes gasperettii*. In purple is indicated the gene exclusively found in *Cerastes gasperettii*. C) Microsynteny for PLA2 in *Naja naja*, *Azemiops feae*, *Crotalus adamanteus* and *Cerastes gasperettii*. Non-toxic PLA2 and flanking genes are also shown. D) Phylogeny of the PLA2 gene family, with two non-toxic PLA2 as outgroups. Some samples that did not fit in any category have been removed. For a complete phylogeny see Fig. S7. Note that PLA2-gK is present in the phylogeny but not in the microsynteny, as any of the studied species contains it. E) Microsynteny for SVSPs for *Crotalus adamanteus* and *Cerastes gasperettii*. Flanking genes are also shown. F) Phylogeny for SVSPs with a non-toxic outgroup. For the three different phylogenies the groups that contained toxins from *Cerastes gasperettii* are highlighted in bold.

### PLA_2_

Regarding PLA_2_, we report two tandem repeat venomous genes for *Cerastes gasperettii* within the non-toxic *PLA_2_-g2E* and *PLA_2_-g2F* array (Fig. 4C), flanked by *OTUD3* and *MUL1* non-toxic genes, as previously reported in other species (Dowell et al., 2016; Margres et al., 2021a; Myers et al., 2022). The number of venomous PLA_2_ in *Cerastes gasperettii* was lower than in *A. feae* and *Crotalus adamanteus*. This difference may be expected, as PLA_2_ only represents around 5% of the proteome for *Cerastes gasperettii* (Fig. 3B) whilst PLA2 are abundant toxins in the proteome for the other two species (Margres et al., 2014; Myers et al., 2022). Phylogenetic results for PLA_2_ genes showed a fully supported clade containing both non-toxic *PLA_2_-g2E* and *PLA_2_-g2F* as outgroups (Fig. 4D and Fig. S7). We also found all other PLA_2_ groups reported in previous studies: *PLA_2_-gC*, *PLA_2_-gK*, *PLA_2_-gB*, *PLA_2_-gD* and *PLA_2_-gA* (Dowell et al., 2016; Myers et al., 2022). The two genes for our target species clustered in different groups (Fig. 4D and Fig. S7). The first PLA_2_ was a *PLA_2_-gD*, which is a group of PLA_2_s exclusively found in true vipers (subfamily Viperinae). The second one was a *PLA_2_-gC* which is more ancestral as it is also found in other pitvipers and non-venomous snakes such as pythons (Dowell et al., 2016). The genomic results are consistent with the proteomics, indicating that specific duplications of PLA_2_ toxin genes have not occurred in *Cerastes gasperettii*.

### SVSPs

Finally, we found eight different SVSPs within the genome of *Cerastes gasperettii*, flanked by *RBM42* and *GRAMD1A* non-toxic genes (Fig. 4E). For this toxin family, we were only able to compare the results with *Crotalus adamanteus*. We did not determine with enough confidence the location of SVSPs within the *N. naja* genome (several regions were matching our venomous SVSP genes as well as the flanking genes). Moreover, *A. feae* was also discarded as SVSPs were not assembled in a single contig. The high number of SVSP genes found (although lower than in *Crotalus adamanteus*) were in line with the proteomic results, as SVSPs are the most abundant toxin in the proteome (Fig. 3B). Phylogenetic results showed three clades, with two of them containing *Cerastes gasperettii* genes (Fig. 4F and Fig. S8). Group 1 was mainly present within *Crotalus*, although there was the presence of some true vipers species, but not in *Cerastes gasperettii* (Fig. S8). Group 2 contained six genes within *Crotalus adamanteus* and only two for *Cerastes gasperettii*. Interestingly, Group 3 was expanded in *Cerastes gasperettii* (Fig. 4E) with a total of six copies, while four were found within *Crotalus adamanteus*. Most of the toxin peptides included in the analyses for true vipers were also found in Group 3 (Fig. S8), indicating a possible expansion of this group of toxins in true vipers (or gene losses in pit vipers). Overall, our high-quality chromosome level reference genome has shed light on the evolution of the main toxin gene families, indicating a compelling correlation between the abundance of toxin genes and the prevalence of these toxins in the venom of *Cerastes gasperettii*.

### Genomic diversity and ancient demographic history

The Arabian horned viper (*C. gasperettii*) is a widespread species, categorized as Least Concern by the IUCN (Egan et al., 2012). Genome-wide diversity was in line with its conservation status, as it showed similar heterozygosity levels compared to other venomous snakes (Fig. 5A). However, more individuals should be sampled along its distribution to verify that similar heterozygosity levels are found across its range. PSMC analyses showed several population expansions and contractions in the last 400 kya, whilst the effective population size of *Cerastes gasperettii* remained relatively constant from 1 until 10 Mya (Fig. 5B). Interestingly, population expansions were coincident with the Last glacial and Penultimate glacial periods (grey lines on Fig. 5B), with a large population increase during the Penultimate Glacial Period (PGP) (1.94×10^5^ to 1.35×10^5^ mya) (Fig. 5B). In fact, during glacial periods, global sea level dropped around 150 meters, exposing the floor and the sand to the wind, which promoted aridification in the Arabian Peninsula and potentially increased habitat suitability for the species (Burriel-Carranza et al., 2023; Glennie & Singhvi, 2002).

**Fig. 5:**
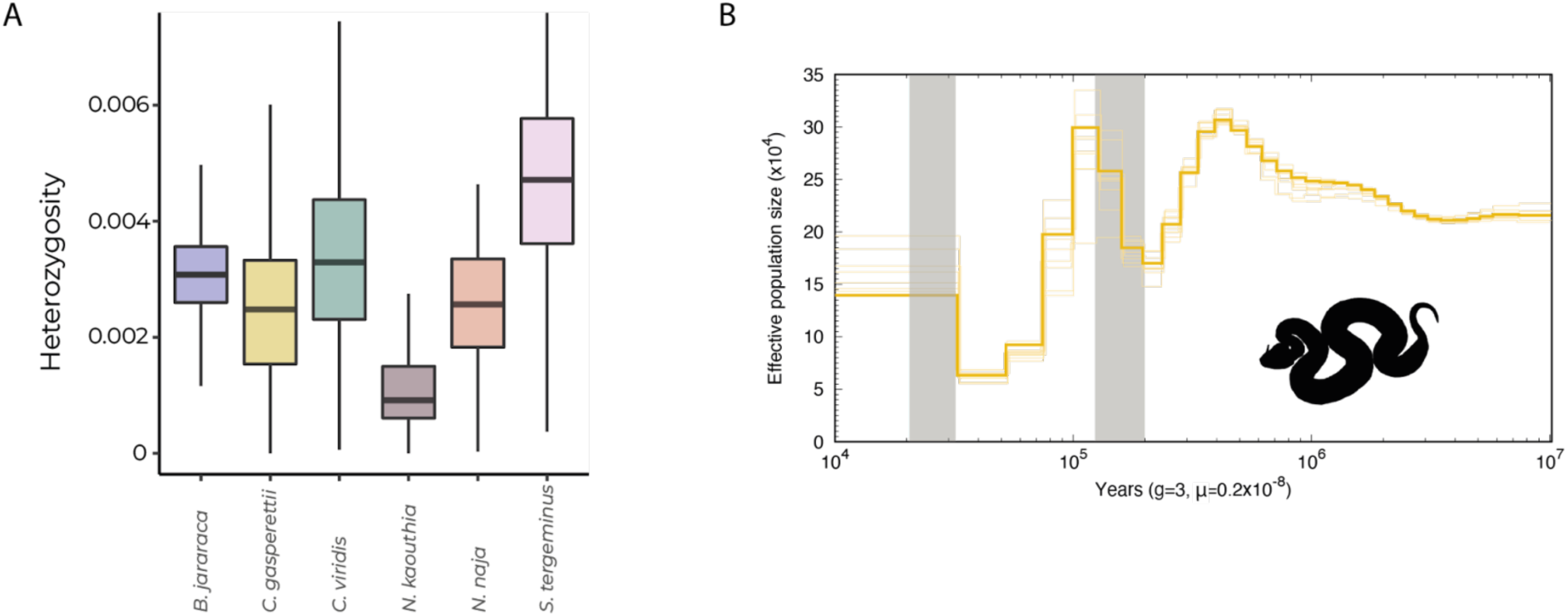
A) Genome-wide diversity for a total of six different venomous snakes: *Bothrops jararaca*, *Cerastes gasperettii*, *Crotalus viridis*, *Naja kaouthia*, *Naja naja* and *Sistrurus tergeminus*. B) PSMC analysis recovering the ancient demographic history of *Cerastes gasperettii*. Generation time was set to 3 years and the substitution rate to 2.4×10^-9^ per site per year. Shaded lines represent 10 bootstrap estimates. Two last glacial periods are shown with grey lines.

## Conclusions

Our high-quality chromosome-level reference genome showed that chromosomal architecture is highly conserved between Crotalinae and Viperinae subfamilies, and differs from elapid genomes by a small number of chromosomal rearrangements. We also found the genomic coordinates of the main toxin-encoding genes, highlighting gene duplication as the main driver in the evolution of SVMP and SVSP toxins. We identified a new SVMP toxin gene, showcasing the importance of using high-quality reference genomes (combined with other -omic techniques) for thoroughly characterizing toxin-encoding genes. Finally, this is a new and important resource for a large clade with currently few reference genomes available. Future genomic studies focusing on Old World viper evolution will benefit greatly from this resource, which will help unveil the origin and diversification of venom and serve as an essential genomic tool for further venomic studies on the subfamily Viperinae.

## Acknowledgements

GM-R is supported by an FPI grant from the Ministerio de Ciencia, Innovación y Universidades, Spain (PRE2019-088729), SRH is awarded by the National Science Foundation Graduate Research Fellowship Program with grant no. 2136515, AT is supported by “la Caixa” doctoral fellowship program (LCF/BQ/DR20/11790007), BB-C is supported by FPU grant from Ministerio de Ciencia, Innovación y Universidades, Spain (FPU18/04742) and ME is supported by an FPI grant from Ministerio de Ciencia e Innovación (PRE2022-101473). In the UAE, we wish to thank His Highness Sheikh Dr. Sultan bin Mohammed Al Qasimi, Supreme Council Member and Ruler of Sharjah, H. E. Ms. Hana Saif al Suwaidi (Chairperson of the Environment and Protected Areas Authority, Sharjah) for their continuous support. We thank Jonathan Wood and Klara Eleftheriadi for their input during the genome assembly and manual curation processes. We also thank Valéria Marques for her help in building the figures and Prem Aguilar for reviewing a previous version of the manuscript.

## Data availability

Final assembly as well as the annotation file were deposited in NCBI under bioproject No. XXX. The final assembly can be accessed with XXX; PacBio subreads with SRRXXX and RNA-seq raw reads with XXX.

## Funding

This work was funded by grant PID2021-128901NB-I00 (MCIN/AEI/10.13039/501100011033 and by ERDF, A way of making Europe; Spain) and grant 2021-SGR-00751 from the Departament de Recerca i Universitats from the Generalitat de Catalunya, Spain to SC.

## Competing Interests

The authors declare that they have no competing interests.

## Author’s contribution

Conceptualization: G.M.R., A.T., B.B.C., J.C., J.E., M.M., S.C. Investigation: S.H, V.P., M.E., T.B., S.B., M.H., J.T.G., D.P., J.C., M.M. Funding acquisition: S.C. Writing-original draft: G.M.R. Writing-review & editing: All authors read, revised, and approved the manuscript final version.

## Ethics statement

No in vivo experiments were performed. Specimens were collected and manipulated with the authorization and under strict control and permission of the government of the United Arab Emirates (Environment and Protected Areas Authority, Government of Sharjah), who approved the study. Specimens were captured and processed following the guidelines and protocols stated in the agreements obtained from the competent authority of the United Arab Emirates. Members of the government supervised collecting activities. All efforts were made to minimize animal suffering. All the research in the United Arab Emirates was done under the supervision and permission of the Environment and Protected Areas Authority, Government of Sharjah.

## Notes

### Competing Interest Statement

The authors have declared no competing interest.

## References

1. Alföldi, J., Di Palma, F., Grabherr, M., Williams, C., Kong, L., Mauceli, E., Russell, P., Lowe, C. B., Glor, R. E., Jaffe, J. D., Ray, D. A., Boissinot, S., Shedlock, A. M., Botka, C., Castoe, T. A., Colbourne, J. K., Fujita, M. K., Moreno, R. G., Ten Hallers, B. F., … Lindblad-Toh, K. (2011). The genome of the green anole lizard and a comparative analysis with birds and mammals. Nature, 477(7366), 587–591. 10.1038/nature10390

2. Ali, S. A., Jackson, T. N. W., Casewell, N. R., Low, D. H. W., Rossi, S., Baumann, K., Fathinia, B., Visser, J., Nouwens, A., Hendrikx, I., Jones, A., Undheim, E. A., & Fry, B. G. (2015). Extreme venom variation in Middle Eastern vipers: A proteomics comparison of Eristicophis macmahonii, Pseudocerastes fieldi and Pseudocerastes persicus. Journal of Proteomics, 116, 106–113. 10.1016/j.jprot.2014.09.003

3. Allio, R., Schomaker-Bastos, A., Romiguier, J., Prosdocimi, F., Nabholz, B., & Delsuc, F. (2020). MitoFinder: Efficient automated large-scale extraction of mitogenomic data in target enrichment phylogenomics. Molecular Ecology Resources, 20(4), 892–905. 10.1111/1755-0998.13160

4. Almeida, D. D., Viala, V. L., Nachtigall, P. G., Broe, M., Gibbs, H. L., Serrano, S. M. D. T., Moura-da-Silva, A. M., Ho, P. L., Nishiyama-Jr, M. Y., & Junqueira-de-Azevedo, I. L. M. (2021). Tracking the recruitment and evolution of snake toxins using the evolutionary context provided by the *Bothrops jararaca* genome. Proceedings of the National Academy of Sciences, 118(20), e2015159118. 10.1073/pnas.2015159118

5. Al-Sadoon, M. K., & Paray, B. A. (2016). Ecological aspects of the horned viper, Cerastes cerastes gasperettii in the central region of Saudi Arabia. Saudi Journal of Biological Sciences, 23(1), 135–138. 10.1016/j.sjbs.2015.10.010

6. Amr, Z. S., Abu Baker, M. A., & Warrell, D. A. (2020). Terrestrial venomous snakes and snakebites in the Arab countries of the Middle East. Toxicon, 177, 1–15. 10.1016/j.toxicon.2020.01.012

7. Andrews, S. (2010). FastQC: a quality control tool for high throughput sequence data.

8. Arnold, N. E., Robinson, M. D., & Carranza, S. (2009). A preliminary analysis of phylogenetic relationships and biogeography of the dangerously venomous Carpet Vipers, Echis (Squamata, Serpentes, Viperidae) based on mitochondrial DNA sequences. Amphibia Reptilia, 30(2), 273–282. 10.1163/156853809788201090

9. Avella, I., Calvete, J. J., Sanz, L., Wüster, W., Licata, F., Quesada-Bernat, S., Rodríguez, Y., & Martínez-Freiría, F. (2022). Interpopulational variation and ontogenetic shift in the venom composition of Lataste’s viper (Vipera latastei, Boscá 1878) from northern Portugal. *Journal of Proteomics*, *263*, 104613. 10.1016/j.jprot.2022.104613

10. Bao, W., Kojima, K. K., & Kohany, O. (2015). Repbase Update, a database of repetitive elements in eukaryotic genomes. Mobile DNA, 6(1), 11. 10.1186/s13100-015-0041-9

11. Broad Institute. (2021). *Picard Tools*. Broad Institute, GitHub Repository.

12. Burriel-Carranza, B., Tejero-Cicuéndez, H., Carné, A., Riaño, G., Talavera, A., Saadi, S. A., Els, J., Šmíd, J., Tamar, K., Tarroso, P., & Carranza, S. (2023). The origin of a mountain biota: Hyper-aridity shaped reptile diversity in an Arabian biodiversity hotspot. 10.1101/2023.04.07.536010

13. Bylsma, R., Walkup, D. K., Hibbitts, T. J., Ryberg, W. A., Black, A. N., & DeWoody, J. A. (2022). Population genetic and genomic analyses of Western Massasauga (Sistrurus tergeminus ssp.): Implications for subspecies delimitation and conservation. Conservation Genetics, 23(2), 271–283. 10.1007/s10592-021-01420-8

14. Calvete, J. J., Pla, D., Els, J., Carranza, S., Damm, M., Hempel, B.-F., John, E. B. O., Petras, D., Heiss, P., Nalbantsoy, A., Göçmen, B., Süssmuth, R. D., Calderón-Celis, F., Nosti, A. J., & Encinar, J. R. (2021). Combined Molecular and Elemental Mass Spectrometry Approaches for Absolute Quantification of Proteomes: Application to the Venomics Characterization of the Two Species of Desert Black Cobras, *Walterinnesia aegyptia* and *Walterinnesia morgani*. Journal of Proteome Research, 20(11), 5064–5078. 10.1021/acs.jproteome.1c00608

15. Carranza, S., Els, J., & Burriel-Carranza, B. (2021). *A field guide to the reptiles of Oman*.

16. Casewell, N. R., Harrison, R. A., Wüster, W., & Wagstaff, S. C. (2009). Comparative venom gland transcriptome surveys of the saw-scaled vipers (Viperidae: Echis) reveal substantial intra-family gene diversity and novel venom transcripts. BMC Genomics, 10(1), 564. 10.1186/1471-2164-10-564

17. Casewell, N. R., Wagstaff, S. C., Wüster, W., Cook, D. A. N., Bolton, F. M. S., King, S. I., Pla, D., Sanz, L., Calvete, J. J., & Harrison, R. A. (2014). Medically important differences in snake venom composition are dictated by distinct postgenomic mechanisms. Proceedings of the National Academy of Sciences, 111(25), 9205–9210. 10.1073/pnas.1405484111

18. Casewell, N. R., Wüster, W., Vonk, F. J., Harrison, R. A., & Fry, B. G. (2013). Complex cocktails: The evolutionary novelty of venoms. Trends in Ecology & Evolution, 28(4), 219–229. 10.1016/j.tree.2012.10.020

19. Challis, R., Richards, E., Rajan, J., Cochrane, G., & Blaxter, M. (2020). BlobToolKit – Interactive Quality Assessment of Genome Assemblies. G3 Genes|Genomes|Genetics, 10(4), 1361–1374. 10.1534/g3.119.400908

20. Chen, S., Zhou, Y., Chen, Y., & Gu, J. (2018). Fastp: An ultra-fast all-in-one FASTQ preprocessor. Bioinformatics, 34(17), i884–i890. 10.1093/bioinformatics/bty560

21. Cheng, H., Concepcion, G. T., Feng, X., Zhang, H., & Li, H. (2021). Haplotype-resolved de novo assembly using phased assembly graphs with hifiasm. Nature Methods, 18(2), Article 2. 10.1038/s41592-020-01056-5

22. Dainat, J., Hereñú, D., Dr. K. D. Murray, Davis, E., Crouch, K., LucileSol, Agostinho, N., Pascal-Git, Zollman, Z., & Tayyrov. (2023). *NBISweden/AGAT: AGAT-v1.2.0* (v1.2.0) [Computer software]. Zenodo. 10.5281/ZENODO.3552717

23. Danecek, P., Bonfield, J. K., Liddle, J., Marshall, J., Ohan, V., Pollard, M. O., Whitwham, A., Keane, T., McCarthy, S. A., Davies, R. M., & Li, H. (2021). Twelve years of SAMtools and BCFtools. GigaScience, 10(2). 10.1093/gigascience/giab008

24. Dowell, N. L., Giorgianni, M. W., Kassner, V. A., Selegue, J. E., Sanchez, E. E., & Carroll, S. B. (2016). The Deep Origin and Recent Loss of Venom Toxin Genes in Rattlesnakes. Current Biology, 26(18), 2434– 2445. 10.1016/j.cub.2016.07.038

25. Drukewitz, S. H., & Von Reumont, B. M. (2019). The Significance of Comparative Genomics in Modern Evolutionary Venomics. Frontiers in Ecology and Evolution, 7, 163. 10.3389/fevo.2019.00163

26. Dussex, N., van der Valk, T., Morales, H. E., Wheat, C. W., Díez-del-Molino, D., von Seth, J., Foster, Y., Kutschera, V. E., Guschanski, K., Rhie, A., Phillippy, A. M., Korlach, J., Howe, K., Chow, W., Pelan, S., Mendes Damas, J. D., Lewin, H. A., Hastie, A. R., Formenti, G., … Dalén, L. (2021). Population genomics of the critically endangered kākāpō. Cell Genomics, 1(1), 100002. 10.1016/j.xgen.2021.100002

27. Egan, D., Amr, Z., Al Johany, A., Els, J., Papenfuss, T., Nilson Sadek, R., Disi, A., Hraoui-Bloquet, S., Werner, Y., & Anderson, S. (2012). *The IUCN Red List of Threatened Species: Cerastes gasperettii* [dataset]. 10.2305/IUCN.UK.2012.RLTS.T164599A1060588.en

28. Fahmi, L., Makran, B., Pla, D., Sanz, L., Oukkache, N., Lkhider, M., Harrison, R. A., Ghalim, N., & Calvete, J. J. (2012). Venomics and antivenomics profiles of North African Cerastes cerastes and C. vipera populations reveals a potentially important therapeutic weakness. Journal of Proteomics, 75(8), 2442– 2453. 10.1016/j.jprot.2012.02.021

29. Ferraz, C. R., Arrahman, A., Xie, C., Casewell, N. R., Lewis, R. J., Kool, J., & Cardoso, F. C. (2019). Multifunctional Toxins in Snake Venoms and Therapeutic Implications: From Pain to Hemorrhage and Necrosis. Frontiers in Ecology and Evolution, 7. https://www.frontiersin.org/articles/10.3389/fevo.2019.00218

30. Flynn, J. M., Hubley, R., Goubert, C., Rosen, J., Clark, A. G., Feschotte, C., & Smit, A. F. (2020). RepeatModeler2 for automated genomic discovery of transposable element families. Proceedings of the National Academy of Sciences, 117(17), 9451–9457. 10.1073/pnas.1921046117

31. Formenti, G., Abueg, L., Brajuka, A., Brajuka, N., Gallardo-Alba, C., Giani, A., Fedrigo, O., & Jarvis, E. D. (2022). Gfastats: Conversion, evaluation and manipulation of genome sequences using assembly graphs. Bioinformatics, 38(17), 4214–4216. 10.1093/bioinformatics/btac460

32. Frantz, L. A. F., Bradley, D. G., Larson, G., & Orlando, L. (2020). Animal domestication in the era of ancient genomics. Nature Reviews Genetics, 21(8), Article 8. 10.1038/s41576-020-0225-0

33. Fry, B. (Ed.). (2015). Venomous reptiles and their toxins: Evolution, pathophysiology, and biodiscovery. Oxford University Press.

34. Fry, B. G., Roelants, K., Champagne, D. E., Scheib, H., Tyndall, J. D. A., King, G. F., Nevalainen, T. J., Norman, J. A., Lewis, R. J., Norton, R. S., Renjifo, C., & de la Vega, R. C. R. (2009). The toxicogenomic multiverse: Convergent recruitment of proteins into animal venoms. Annual Review of Genomics and Human Genetics, 10, 483–511. 10.1146/annurev.genom.9.081307.164356

35. Fry, B. G., Scheib, H., van der Weerd, L., Young, B., McNaughtan, J., Ramjan, S. F. R., Vidal, N., Poelmann, R. E., & Norman, J. A. (2008). Evolution of an Arsenal: Structural and Functional Diversification of the Venom System in the Advanced Snakes (Caenophidia)*. Molecular & Cellular Proteomics, 7(2), 215–246. 10.1074/mcp.M700094-MCP200

36. Fry, B. G., & Wüster, W. (2004). Assembling an Arsenal: Origin and Evolution of the Snake Venom Proteome Inferred from Phylogenetic Analysis of Toxin Sequences. Molecular Biology and Evolution, 21(5), 870–883. 10.1093/molbev/msh091

37. Gabriel, L., Hoff, K. J., Brůna, T., Borodovsky, M., & Stanke, M. (2021). TSEBRA: Transcript selector for BRAKER. BMC Bioinformatics, 22(1), 566. 10.1186/s12859-021-04482-0

38. Geneva, A. J., Park, S., Bock, D. G., De Mello, P. L. H., Sarigol, F., Tollis, M., Donihue, C. M., Reynolds, R. G., Feiner, N., Rasys, A. M., Lauderdale, J. D., Minchey, S. G., Alcala, A. J., Infante, C. R., Kolbe, J. J., Schluter, D., Menke, D. B., & Losos, J. B. (2022). Chromosome-scale genome assembly of the brown anole (Anolis sagrei), an emerging model species. Communications Biology, 5(1), 1126. 10.1038/s42003-022-04074-5

39. Ghurye, J., Rhie, A., Walenz, B. P., Schmitt, A., Selvaraj, S., Pop, M., Phillippy, A. M., & Koren, S. (2019). Integrating Hi-C links with assembly graphs for chromosome-scale assembly. PLoS Computational Biology, 15(8), e1007273. 10.1371/journal.pcbi.1007273

40. Gilbert, C., Meik, J. M., Dashevsky, D., Card, D. C., Castoe, T. A., & Schaack, S. (2014). Endogenous hepadnaviruses, bornaviruses and circoviruses in snakes. Proceedings of the Royal Society B: Biological Sciences, 281(1791), 20141122. 10.1098/rspb.2014.1122

41. Giorgianni, M. W., Dowell, N. L., Griffin, S., Kassner, V. A., Selegue, J. E., & Carroll, S. B. (2020). The origin and diversification of a novel protein family in venomous snakes. Proceedings of the National Academy of Sciences, 117(20), 10911–10920. 10.1073/pnas.1920011117

42. Glennie, K. W., & Singhvi, A. K. (2002). Event stratigraphy, paleoenvironment and chronology of SE Arabian deserts. Quaternary Science Reviews, 21(7), 853–869. 10.1016/S0277-3791(01)00133-0

43. Green, R. E., Braun, E. L., Armstrong, J., Earl, D., Nguyen, N., Hickey, G., Vandewege, M. W., St. John, J. A., Capella-Gutiérrez, S., Castoe, T. A., Kern, C., Fujita, M. K., Opazo, J. C., Jurka, J., Kojima, K. K., Caballero, J., Hubley, R. M., Smit, A. F., Platt, R. N., … Ray, D. A. (2014). Three crocodilian genomes reveal ancestral patterns of evolution among archosaurs. Science, 346(6215), 1254449. 10.1126/science.1254449

44. Guan, D., McCarthy, S. A., Wood, J., Howe, K., Wang, Y., & Durbin, R. (2020). Identifying and removing haplotypic duplication in primary genome assemblies. Bioinformatics, 36(9), 2896–2898. 10.1093/bioinformatics/btaa025

45. Guindon, S., Dufayard, J.-F., Lefort, V., Anisimova, M., Hordijk, W., & Gascuel, O. (2010). New Algorithms and Methods to Estimate Maximum-Likelihood Phylogenies: Assessing the Performance of PhyML 3.0. Systematic Biology, 59(3), 307–321. 10.1093/sysbio/syq010

46. Gurevich, A., Saveliev, V., Vyahhi, N., & Tesler, G. (2013). QUAST: Quality assessment tool for genome assemblies. *Bioinformatics (Oxford*, England), 29(8), 1072–1075. 10.1093/bioinformatics/btt086

47. Gutiérrez, J. M., Calvete, J. J., Habib, A. G., Harrison, R. A., Williams, D. J., & Warrell, D. A. (2017). Snakebite envenoming. Nature Reviews Disease Primers, 3(1), Article 1. 10.1038/nrdp.2017.63

48. Hogan, M. P., Holding, M. L., Nystrom, G. S., Colston, T. J., Bartlett, D. A., Mason, A. J., Ellsworth, S. A., Rautsaw, R. M., Lawrence, K. C., Strickland, J. L., He, B., Fraser, P., Margres, M. J., Gilbert, D. M., Gibbs, H. L., Parkinson, C. L., & Rokyta, D. R. (2024). The genetic regulatory architecture and epigenomic basis for age-related changes in rattlesnake venom. Proceedings of the National Academy of Sciences, 121(16), e2313440121. 10.1073/pnas.2313440121

49. Hogan, M. P., Whittington, A. C., Broe, M. B., Ward, M. J., Gibbs, H. L., & Rokyta, D. R. (2021). The Chemosensory Repertoire of the Eastern Diamondback Rattlesnake (Crotalus adamanteus) Reveals Complementary Genetics of Olfactory and Vomeronasal-Type Receptors. Journal of Molecular Evolution, 89(4), 313–328. 10.1007/s00239-021-10007-3

50. Jan, V., Maroun, R. C., Robbe-Vincent, A., De Haro, L., & Choumet, V. (2002). Toxicity evolution of Vipera aspis aspis venom: Identification and molecular modeling of a novel phospholipase A2 heterodimer neurotoxin11Nucleotide sequence data reported are available in the EMBL database under the accession numbers AJ459806 and AJ459807. FEBS Letters, 527(1), 263–268. 10.1016/S0014-5793(02)03205-2

51. Jin, J.-J., Yu, W.-B., Yang, J.-B., Song, Y., dePamphilis, C. W., Yi, T.-S., & Li, D.-Z. (2020). GetOrganelle: A fast and versatile toolkit for accurate de novo assembly of organelle genomes. Genome Biology, 21(1), 241. 10.1186/s13059-020-02154-5

52. Jones, P., Binns, D., Chang, H.-Y., Fraser, M., Li, W., McAnulla, C., McWilliam, H., Maslen, J., Mitchell, A., Nuka, G., Pesseat, S., Quinn, A. F., Sangrador-Vegas, A., Scheremetjew, M., Yong, S.-Y., Lopez, R., & Hunter, S. (2014). InterProScan 5: Genome-scale protein function classification. Bioinformatics, 30(9), 1236–1240. 10.1093/bioinformatics/btu031

53. Kalita, B., Mackessy, S. P., & Mukherjee, A. K. (2018). Proteomic analysis reveals geographic variation in venom composition of Russell’s Viper in the Indian subcontinent: Implications for clinical manifestations post-envenomation and antivenom treatment. Expert Review of Proteomics, 15(10), 837–849. 10.1080/14789450.2018.1528150

54. Katoh, K., & Standley, D. M. (2013). MAFFT Multiple Sequence Alignment Software Version 7: Improvements in Performance and Usability. Molecular Biology and Evolution, 30(4), 772–780. 10.1093/molbev/mst010

55. Keilwagen, J., Hartung, F., & Grau, J. (2019). GeMoMa: Homology-Based Gene Prediction Utilizing Intron Position Conservation and RNA-seq Data. In M. Kollmar (Ed.), Gene Prediction (Vol. 1962, pp. 161– 177). Springer New York. 10.1007/978-1-4939-9173-0_9

56. Kiełbasa, S. M., Wan, R., Sato, K., Horton, P., & Frith, M. C. (2011). Adaptive seeds tame genomic sequence comparison. Genome Research, 21(3), 487–493. 10.1101/gr.113985.110

57. Kim, D., Paggi, J. M., Park, C., Bennett, C., & Salzberg, S. L. (2019). Graph-based genome alignment and genotyping with HISAT2 and HISAT-genotype. Nature Biotechnology, 37(8), 907–915. 10.1038/s41587-019-0201-4

58. King, G. F. (2011). Venoms as a platform for human drugs: Translating toxins into therapeutics. Expert Opinion on Biological Therapy, 11(11), 1469–1484. 10.1517/14712598.2011.621940

59. Li, D., Luo, R., Liu, C.-M., Leung, C.-M., Ting, H.-F., Sadakane, K., Yamashita, H., & Lam, T.-W. (2016). MEGAHIT v1.0: A fast and scalable metagenome assembler driven by advanced methodologies and community practices. Methods, 102, 3–11. 10.1016/j.ymeth.2016.02.020

60. Li, H. (2013). Aligning sequence reads, clone sequences and assembly contigs with BWA-MEM. http://arxiv.org/abs/1303.3997

61. Li, H., & Durbin, R. (2011). Inference of human population history from individual whole-genome sequences. Nature 2011 475:7357, 475(7357), 493–496. 10.1038/nature10231

62. Li, H., Handsaker, B., Wysoker, A., Fennell, T., Ruan, J., Homer, N., Marth, G., Abecasis, G., & Durbin, R. (2009). The Sequence Alignment/Map format and SAMtools. Bioinformatics, 25(16), 2078–2079. 10.1093/bioinformatics/btp352

63. Li, L., Huang, J., & Lin, Y. (2018). Snake Venoms in Cancer Therapy: Past, Present and Future. Toxins, 10(9), 346. 10.3390/toxins10090346

64. Love, M. I., Huber, W., & Anders, S. (2014). Moderated estimation of fold change and dispersion for RNA-seq data with DESeq2. Genome Biology, 15(12), 550. 10.1186/s13059-014-0550-8

65. Mackessy, S. P. (2010). Evolutionary trends in venom composition in the Western Rattlesnakes (Crotalus viridis sensu lato): Toxicity vs. tenderizers. Toxicon, 55(8), 1463–1474. 10.1016/j.toxicon.2010.02.028

66. Margres, M. J., McGivern, J. J., Wray, K. P., Seavy, M., Calvin, K., & Rokyta, D. R. (2014). Linking the transcriptome and proteome to characterize the venom of the eastern diamondback rattlesnake (Crotalus adamanteus). Journal of Proteomics, 96, 145–158. 10.1016/j.jprot.2013.11.001

67. Margres, M. J., Rautsaw, R. M., Strickland, J. L., Mason, A. J., Schramer, T. D., Hofmann, E. P., Stiers, E., Ellsworth, S. A., Nystrom, G. S., Hogan, M. P., Bartlett, D. A., Colston, T. J., Gilbert, D. M., Rokyta, D. R., & Parkinson, C. L. (2021). The Tiger Rattlesnake genome reveals a complex genotype underlying a simple venom phenotype. Proceedings of the National Academy of Sciences, 118(4), e2014634118. 10.1073/pnas.2014634118

68. Margres, M. J., Wray, K. P., Sanader, D., McDonald, P. J., Trumbull, L. M., Patton, A. H., & Rokyta, D. R. (2021). Varying Intensities of Introgression Obscure Incipient Venom-Associated Speciation in the Timber Rattlesnake (Crotalus horridus). Toxins, 13(11), Article 11. 10.3390/toxins13110782

69. Martin, M. (2011). Cutadapt removes adapter sequences from high-throughput sequencing reads. EMBnet.Journal, 17(1), 10. 10.14806/ej.17.1.200

70. McKenna, A., Hanna, M., Banks, E., Sivachenko, A., Cibulskis, K., Kernytsky, A., Garimella, K., Altshuler, D., Gabriel, S., Daly, M., & DePristo, M. A. (2010). The genome analysis toolkit: A MapReduce framework for analyzing next-generation DNA sequencing data. Genome Research, 20(9), 1297– 1303. 10.1101/gr.107524.110

71. Mochales-Riaño, G., Burriel-Carranza, B., Barros, M. I., Velo-Antón, G., Talavera, A., Spilani, L., Tejero-Cicuéndez, H., Crochet, P.-A., Piris, A., García-Cardenete, L., Busais, S., Els, J., Shobrak, M., Brito, J. C., Šmíd, J., Carranza, S., & Martínez-Freiría, F. (2024). Hidden in the sand: Phylogenomics unravel an unexpected evolutionary history for the desert-adapted vipers of the genus Cerastes. Molecular Phylogenetics and Evolution, 191, 107979. 10.1016/j.ympev.2023.107979

73. Myers, E. A., Strickland, J. L., Rautsaw, R. M., Mason, A. J., Schramer, T. D., Nystrom, G. S., Hogan, M. P., Yooseph, S., Rokyta, D. R., & Parkinson, C. L. (2022). De Novo Genome Assembly Highlights the Role of Lineage-Specific Gene Duplications in the Evolution of Venom in Fea’s Viper (*Azemiops feae*). Genome Biology and Evolution, 14(7), evac082. 10.1093/gbe/evac082

74. Orteu, A., & Jiggins, C. D. (2020). The genomics of coloration provides insights into adaptive evolution. Nature Reviews Genetics, 21(8), Article 8. 10.1038/s41576-020-0234-z

75. Osipov, A., & Utkin, Y. (2023). What Are the Neurotoxins in Hemotoxic Snake Venoms? International Journal of Molecular Sciences, 24(3), Article 3. 10.3390/ijms24032919

76. Pardos-Blas, J. R., Irisarri, I., Abalde, S., Afonso, C. M. L., Tenorio, M. J., & Zardoya, R. (2021). The genome of the venomous snail *Lautoconus ventricosus* sheds light on the origin of conotoxin diversity. GigaScience, 10(5), giab037. 10.1093/gigascience/giab037

77. Pertea, M., Pertea, G. M., Antonescu, C. M., Chang, T.-C., Mendell, J. T., & Salzberg, S. L. (2015). StringTie enables improved reconstruction of a transcriptome from RNA-seq reads. Nature Biotechnology, 33(3), 290–295. 10.1038/nbt.3122

78. Pook, C. E., Joger, U., Stümpel, N., & Wüster, W. (2009). When continents collide: Phylogeny, historical biogeography and systematics of the medically important viper genus Echis (Squamata: Serpentes: Viperidae). Molecular Phylogenetics and Evolution, 53(3), 792–807. 10.1016/j.ympev.2009.08.002

79. R Core Team. (2021a). R: A Language and Environment for Statistical Computing. R Foundation for Statistical Computing. https://www.R-project.org/

80. R Core Team. (2021b). R: A Language and Environment for Statistical Computing. https://www.R-project.org/

81. Ranallo-Benavidez, T. R., Jaron, K. S., & Schatz, M. C. (2020). GenomeScope 2.0 and Smudgeplot for reference-free profiling of polyploid genomes. Nature Communications, 11(1), Article 1. 10.1038/s41467-020-14998-3

82. Rhie, A., McCarthy, S. A., Fedrigo, O., Damas, J., Formenti, G., Koren, S., Uliano-Silva, M., Chow, W., Fungtammasan, A., Gedman, G. L., Cantin, L. J., Thibaud-Nissen, F., Haggerty, L., Lee, C., Ko, B. J., Kim, J., Bista, I., Smith, M., Haase, B., … Jarvis, E. D. (2020). Towards complete and error-free genome assemblies of all vertebrate species (p. 2020.05.22.110833). bioRxiv. 10.1101/2020.05.22.110833

83. Rhie, A., Walenz, B. P., Koren, S., & Phillippy, A. M. (2020). Merqury: Reference-free quality, completeness, and phasing assessment for genome assemblies. Genome Biology, 21(1), 245. 10.1186/s13059-020-02134-9

84. Rokyta, D. R., Margres, M. J., Ward, M. J., & Sanchez, E. E. (2017). The genetics of venom ontogeny in the eastern diamondback rattlesnake (*Crotalus adamanteus*). PeerJ, 5, e3249. 10.7717/peerj.3249

85. Russell, F. E., & Campbell, J. R. (2015). Venomous terrestrial Snakes of the Middle East. Edition Chimaira.

86. Saethang, T., Somparn, P., Payungporn, S., Sriswasdi, S., Yee, K. T., Hodge, K., Knepper, M. A., Chanhome, L., Khow, O., Chaiyabutr, N., Sitprija, V., & Pisitkun, T. (2022). Identification of Daboia siamensis venome using integrated multi-omics data. Scientific Reports, 12(1), Article 1. 10.1038/s41598-022-17300-1

87. San-Jose, L. M., & Roulin, A. (2017). Genomics of coloration in natural animal populations. Philosophical Transactions of the Royal Society B: Biological Sciences, 372(1724), 20160337. 10.1098/rstb.2016.0337

88. Schield, D. R., Card, D. C., Hales, N. R., Perry, B. W., Pasquesi, G. M., Blackmon, H., Adams, R. H., Corbin, A. B., Smith, C. F., Ramesh, B., Demuth, J. P., Betrán, E., Tollis, M., Meik, J. M., Mackessy, S. P., & Castoe, T. A. (2019). The origins and evolution of chromosomes, dosage compensation, and mechanisms underlying venom regulation in snakes. Genome Research, 29(4), 590–601. 10.1101/gr.240952.118

89. Schield, D. R., Perry, B. W., Adams, R. H., Holding, M. L., Nikolakis, Z. L., Gopalan, S. S., Smith, C. F., Parker, J. M., Meik, J. M., DeGiorgio, M., Mackessy, S. P., & Castoe, T. A. (2022). The roles of balancing selection and recombination in the evolution of rattlesnake venom. Nature Ecology & Evolution, 6(9), 1367–1380. 10.1038/s41559-022-01829-5

90. Schneemann, M., Cathomas, R., Laidlaw, S. T., El Nahas, A. M., Theakston, R. D. G., & Warrell, D. A. (2004). Life-threatening envenoming by the Saharan horned viper (Cerastes cerastes) causing micro-angiopathic haemolysis, coagulopathy and acute renal failure: Clinical cases and review. QJM: An International Journal of Medicine, 97(11), 717–727. 10.1093/qjmed/hch118

91. Simão, F. A., Waterhouse, R. M., Ioannidis, P., Kriventseva, E. V., & Zdobnov, E. M. (2015). BUSCO: Assessing genome assembly and annotation completeness with single-copy orthologs. *Bioinformatics (Oxford*, England), 31(19), 3210–3212. 10.1093/bioinformatics/btv351

92. Šmíd, J., & Tolley, K. A. (2019). Calibrating the tree of vipers under the fossilized birth-death model. Scientific Reports, 9(1), 5510. 10.1038/s41598-019-41290-2

93. Smith, C. F., Nikolakis, Z. L., Perry, B. W., Schield, D. R., Meik, J. M., Saviola, A. J., Castoe, T. A., Parker, J., & Mackessy, S. P. (2023). The best of both worlds? Rattlesnake hybrid zones generate complex combinations of divergent venom phenotypes that retain high toxicity. Biochimie. 10.1016/j.biochi.2023.07.008

94. Solovyev, V., Kosarev, P., Seledsov, I., & Vorobyev, D. (2006). Automatic annotation of eukaryotic genes, pseudogenes and promoters. Genome Biology, 7(Suppl 1), S10. 10.1186/gb-2006-7-s1-s10

95. Suryamohan, K., Krishnankutty, S. P., Guillory, J., Jevit, M., Schröder, M. S., Wu, M., Kuriakose, B., Mathew, O. K., Perumal, R. C., Koludarov, I., Goldstein, L. D., Senger, K., Dixon, M. D., Velayutham, D., Vargas, D., Chaudhuri, S., Muraleedharan, M., Goel, R., Chen, Y.-J. J., … Seshagiri, S. (2020). The Indian cobra reference genome and transcriptome enables comprehensive identification of venom toxins. Nature Genetics, 52(1), 106–117. 10.1038/s41588-019-0559-8

96. Tang, H., Bowers, J. E., Wang, X., Ming, R., Alam, M., & Paterson, A. H. (2008). Synteny and Collinearity in Plant Genomes. Science, 320(5875), 486–488. 10.1126/science.1153917

97. Tang, H., Krishnakumar, V., Jingping Li, Tiany, MichelMoser, Maria, & Yim, W. C. (2017). *tanghaibao/jcvi: JCVI v0.7.5* (v0.7.5) [Computer software]. Zenodo. 10.5281/ZENODO.846919

98. Tang, S., Lomsadze, A., & Borodovsky, M. (2015). Identification of protein coding regions in RNA transcripts. Nucleic Acids Research, 43(12), e78–e78. 10.1093/nar/gkv227

99. Tasoulis, T., & Isbister, G. (2017). A Review and Database of Snake Venom Proteomes. Toxins, 9(9), 290. 10.3390/toxins9090290

100. Tempel, S. (2012). Using and Understanding RepeatMasker. In Y. Bigot (Ed.), Mobile Genetic Elements (Vol. 859, pp. 29–51). Humana Press. 10.1007/978-1-61779-603-6_2

101. Thongchum, R., Singchat, W., Laopichienpong, N., Tawichasri, P., Kraichak, E., Prakhongcheep, O., Sillapaprayoon, S., Muangmai, N., Baicharoen, S., Suntrarachun, S., Chanhome, L., Peyachoknagul, S., & Srikulnath, K. (2019). Diversity of PBI-DdeI satellite DNA in snakes correlates with rapid independent evolution and different functional roles. Scientific Reports, 9(1), 15459. 10.1038/s41598-019-51863-w

102. Title, P. O., Singhal, S., Grundler, M. C., Costa, G. C., Pyron, R. A., Colston, T. J., Grundler, M. R., Prates, I., Stepanova, N., Jones, M. E. H., Cavalcanti, L. B. Q., Colli, G. R., Di-Poï, N., Donnellan, S. C., Moritz, C., Mesquita, D. O., Pianka, E. R., Smith, S. A., Vitt, L. J., & Rabosky, D. L. (2024). The macroevolutionary singularity of snakes. Science, 383(6685), 918–923. 10.1126/science.adh2449

103. Uetz, P. (2021). The Reptile Database: Curating the biodiversity literature without funding. Biodiversity Information Science and Standards, 5, e75448. 10.3897/biss.5.75448

104. Vitt, L. J., & Caldwell, J. P. (2014). Herpetology: An introductory biology of amphibians and reptiles (Fourth edition). Elsevier, AP, Academic Press is an imprint of Elsevier.

105. Vonk, F. J., Casewell, N. R., Henkel, C. V., Heimberg, A. M., Jansen, H. J., McCleary, R. J. R., Kerkkamp, H. M. E., Vos, R. A., Guerreiro, I., Calvete, J. J., Wüster, W., Woods, A. E., Logan, J. M., Harrison, R. A., Castoe, T. A., De Koning, A. P. J., Pollock, D. D., Yandell, M., Calderon, D., … Richardson, M. K. (2013). The king cobra genome reveals dynamic gene evolution and adaptation in the snake venom system. Proceedings of the National Academy of Sciences, 110(51), 20651–20656. 10.1073/pnas.1314702110

106. Walker, B. J., Abeel, T., Shea, T., Priest, M., Abouelliel, A., Sakthikumar, S., Cuomo, C. A., Zeng, Q., Wortman, J., Young, S. K., & Earl, A. M. (2014). Pilon: An Integrated Tool for Comprehensive Microbial Variant Detection and Genome Assembly Improvement. PLOS ONE, 9(11), e112963. 10.1371/journal.pone.0112963

107. Weinstein, S. A., White, J., Keyler, D. E., & Warrell, D. A. (2013). Non-front-fanged colubroid snakes: A current evidence-based analysis of medical significance. Toxicon, 69, 103–113. 10.1016/j.toxicon.2013.02.003

108. Werren, J. H., Richards, S., Desjardins, C. A., Niehuis, O., Gadau, J., Colbourne, J. K., THE NASONIA GENOME WORKING GROUP, Beukeboom, L. W., Desplan, C., Elsik, C. G., Grimmelikhuijzen, C. J. P., Kitts, P., Lynch, J. A., Murphy, T., Oliveira, D. C. S. G., Smith, C. D., Zande, L. van de, Worley, K. C., Zdobnov, E. M., … Gibbs, R. A. (2010). Functional and Evolutionary Insights from the Genomes of Three Parasitoid Nasonia Species. Science, 327(5963), 343–348. 10.1126/science.1178028

109. Westeen, E. P., Escalona, M., Holding, M. L., Beraut, E., Fairbairn, C., Marimuthu, M. P. A., Nguyen, O., Perri, R., Fisher, R. N., Toffelmier, E., Shaffer, H. B., & Wang, I. J. (2023). A genome assembly for the southern Pacific rattlesnake, *Crotalus oreganus helleri*, in the western rattlesnake species complex. Journal of Heredity, 114(6), 681–689. 10.1093/jhered/esad045

110. Wickham, H. (2016). ggplot2: Elegant Graphics for Data Analysis. Springer-Verlag New York. https://ggplot2.tidyverse.org

111. Williams, D. J., Faiz, M. A., Abela-Ridder, B., Ainsworth, S., Bulfone, T. C., Nickerson, A. D., Habib, A. G., Junghanss, T., Fan, H. W., Turner, M., Harrison, R. A., & Warrell, D. A. (2019). Strategy for a globally coordinated response to a priority neglected tropical disease: Snakebite envenoming. PLOS Neglected Tropical Diseases, 13(2), e0007059. 10.1371/journal.pntd.0007059

112. Wüster, W., Peppin, L., Pook, C. E., & Walker, D. E. (2008). A nesting of vipers: Phylogeny and historical biogeography of the Viperidae (Squamata: Serpentes). Molecular Phylogenetics and Evolution, 49(2), 445–459. 10.1016/j.ympev.2008.08.019

113. Zancolli, G., Calvete, J. J., Cardwell, M. D., Greene, H. W., Hayes, W. K., Hegarty, M. J., Herrmann, H.-W., Holycross, A. T., Lannutti, D. I., Mulley, J. F., Sanz, L., Travis, Z. D., Whorley, J. R., Wüster, C. E., & Wüster, W. (2019). When one phenotype is not enough: Divergent evolutionary trajectories govern venom variation in a widespread rattlesnake species. Proceedings of the Royal Society B: Biological Sciences, 286(1898), 20182735. 10.1098/rspb.2018.2735

114. Zancolli, G., Reijnders, M., Waterhouse, R. M., & Robinson-Rechavi, M. (2022). Convergent evolution of venom gland transcriptomes across Metazoa. Proceedings of the National Academy of Sciences, 119(1), e2111392119. 10.1073/pnas.2111392119

