## Supplementary information for "Chromosome-level reference genome for the medically important Arabian horned viper (*Cerastes gasperettii*)"

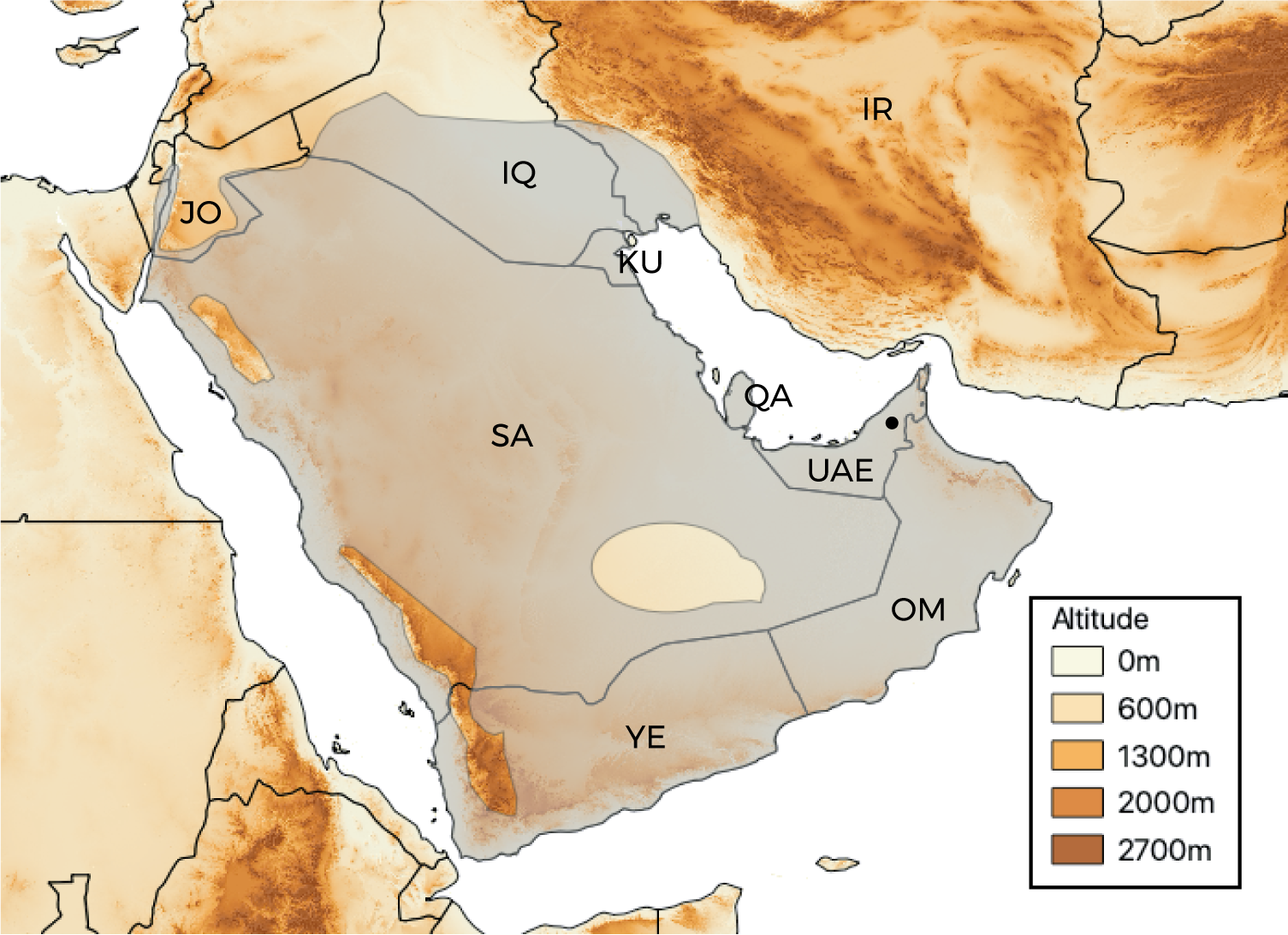

Fig. S1: Distribution map for the studied species *Cerastes gasperettii* with the location of our samples. Countries where the species is present are indicated. Abbreviations are as follows: JO, Jordania; SA, Saudi Arabia; YE, Yemen; OM, Oman; UAE, United Arab Emirates; IQ, Iraq; IR, Iran; KU, Kuwait, QA, Qatar.

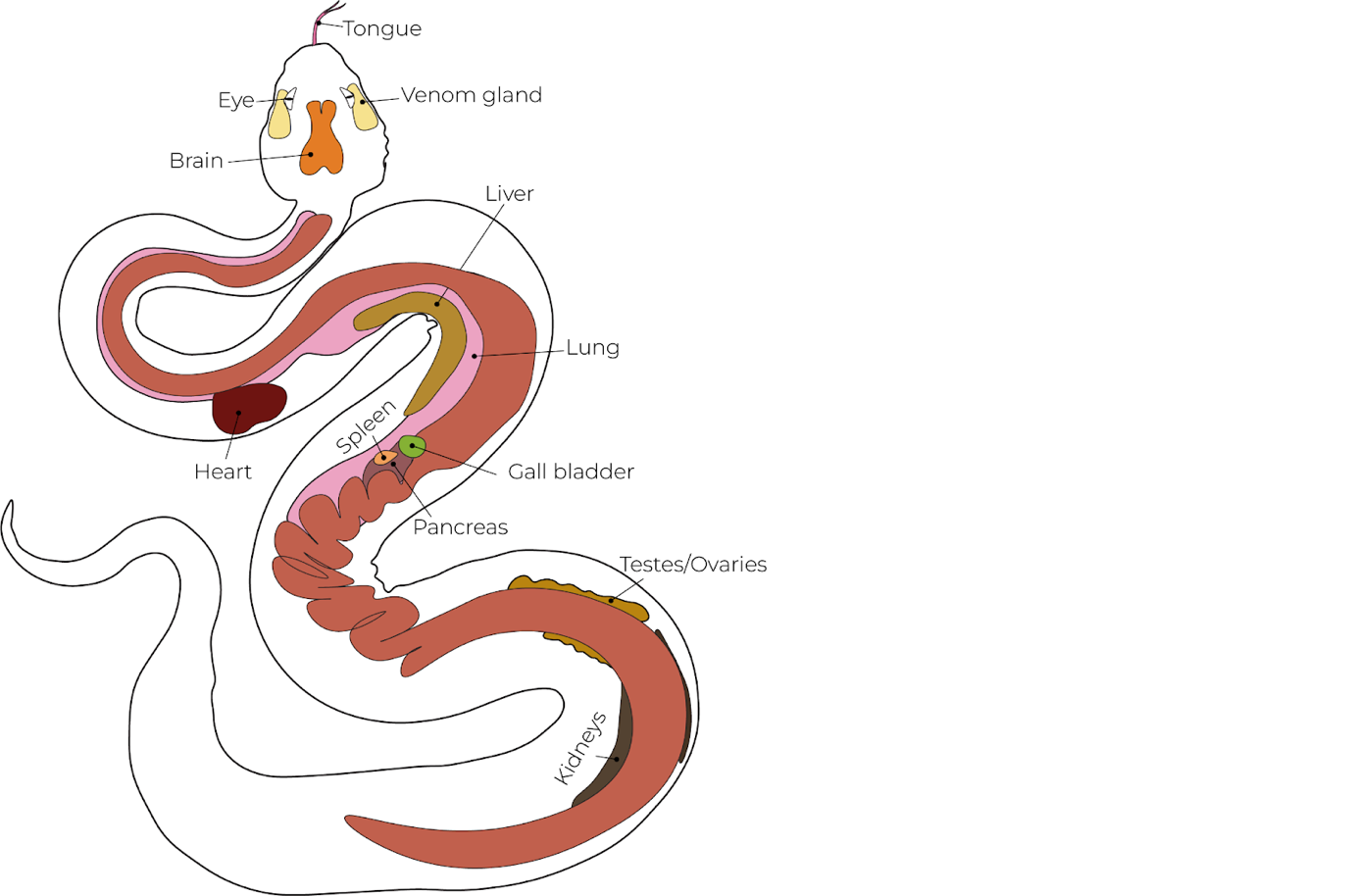

Fig. S2: Drawing of an Arabian horned viper depicting all the tissues sampled for RNA-seq analyses.

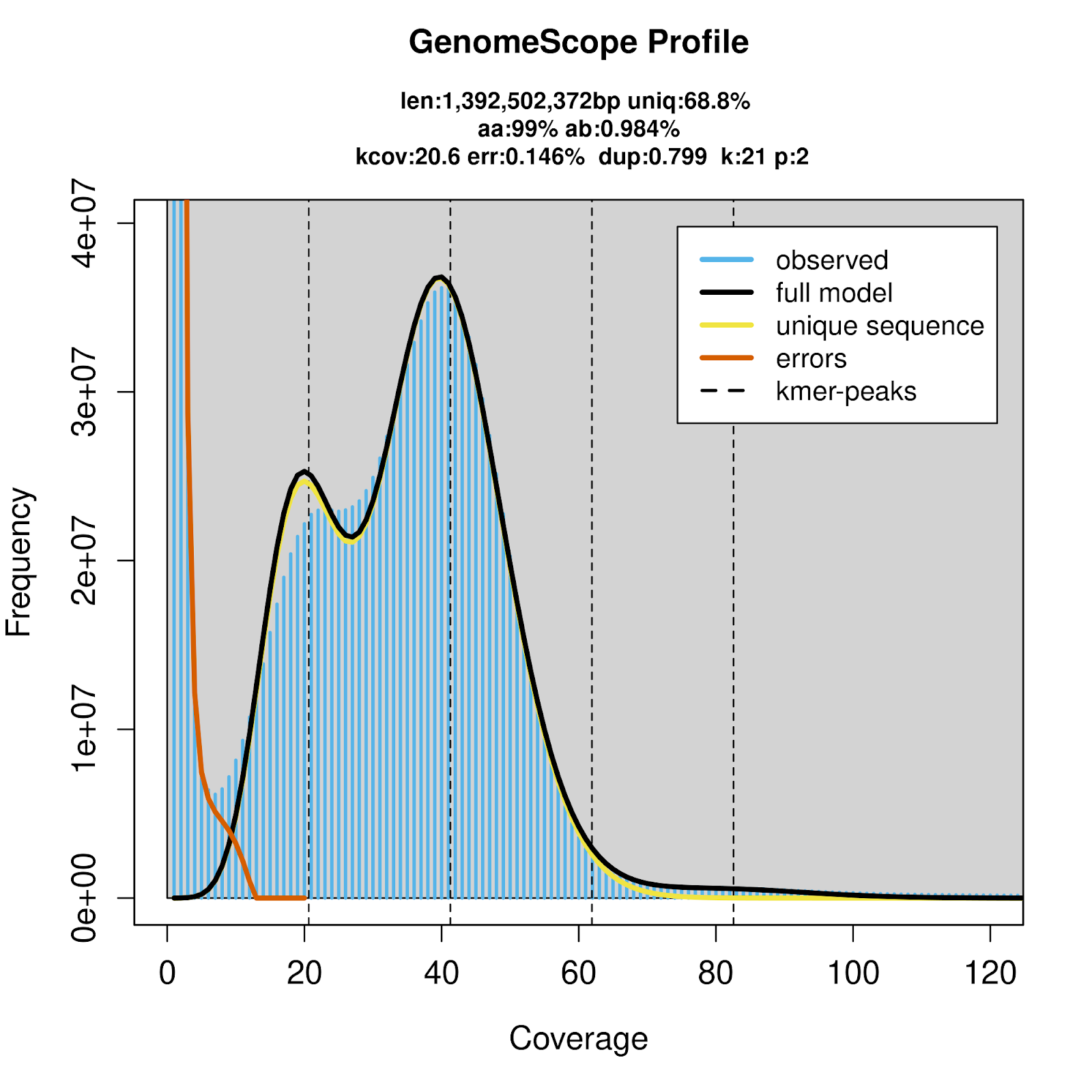

Fig. S3: Histogram from GenomeScope showing the frequency of reads in relation with their coverage.

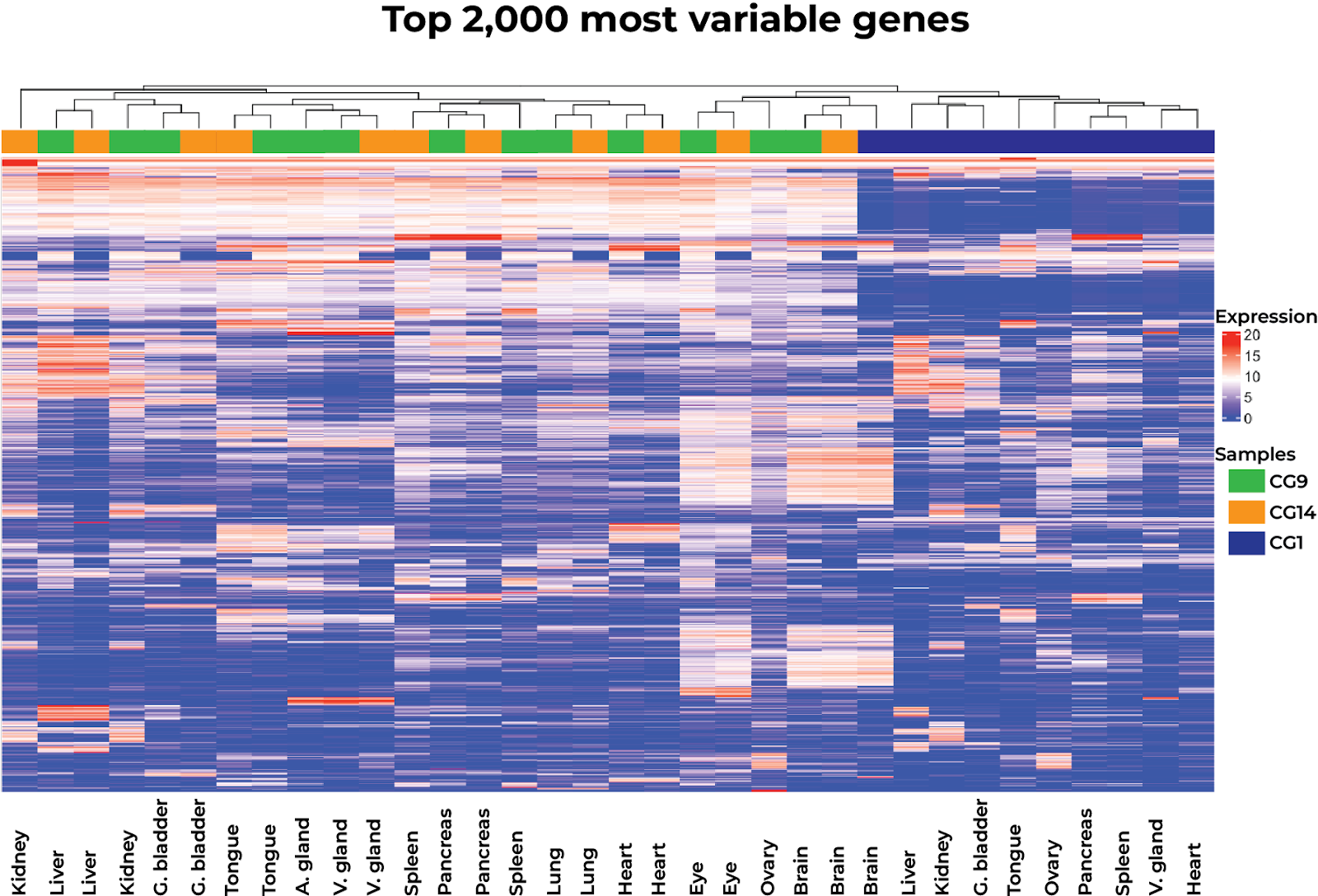

Fig. S4: Heatmap for the 2,000 most variable genes within our three samples, showing a clear batch effect of sample CG1 as well as a high similarity between the putative accessory gland and the venom gland. Each column represents a different sampled tissue. The three different samples are depicted with different colors at the top of the heatmap. Abbreviations are as follows: G. bladder, gallbladder and V. gland, venom gland.

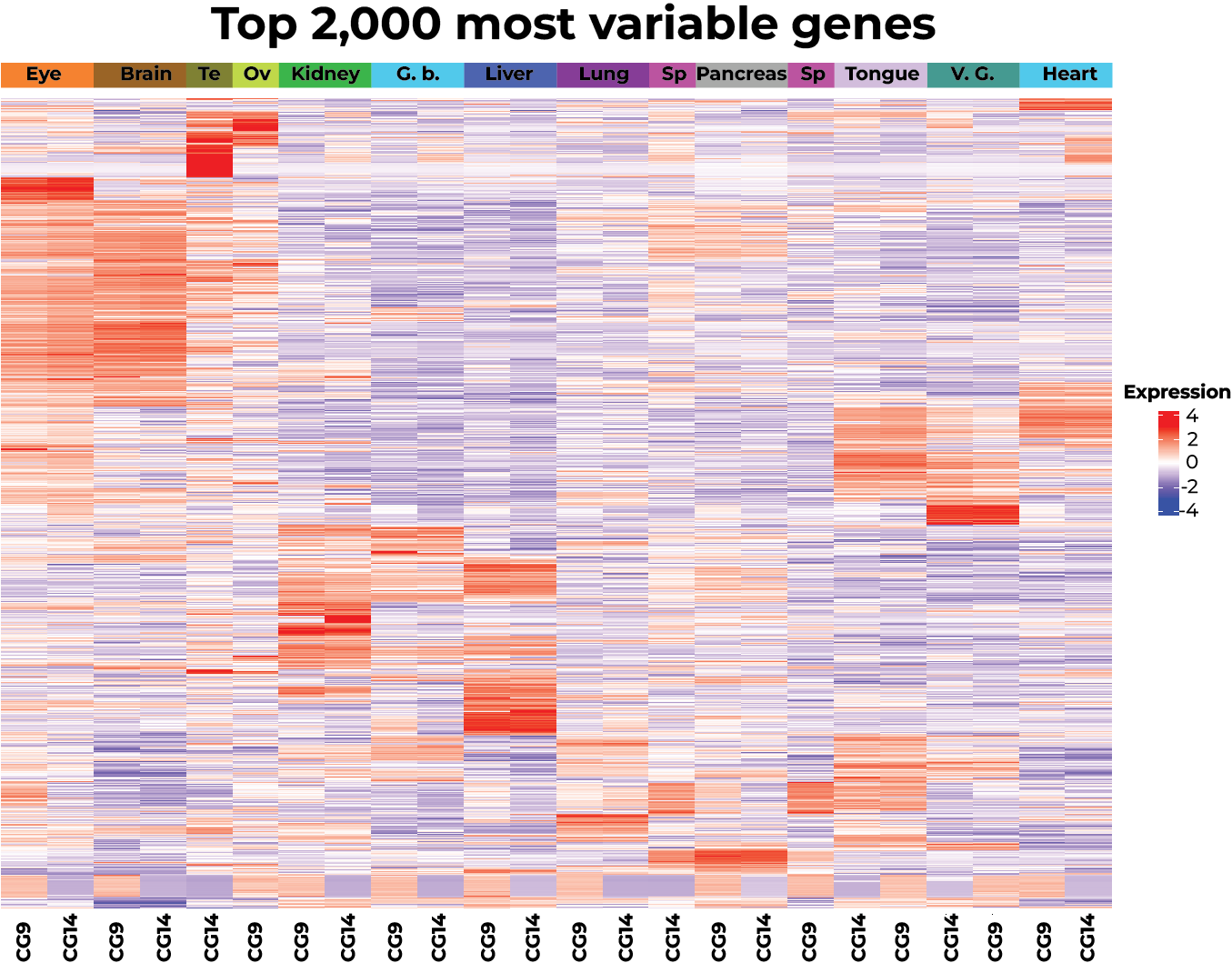

Fig. S5: Heatmap for the 2,000 most variable genes for both samples, reporting highly expressed genes unique for each tissue type. Each column represents one tissue sampled per individual. Expression levels were normalized. Abbreviations are as follows: Te, Testis; Ov, Ovary; G.b., gallbladder; Sp, Spleen and V.G., Venom gland.

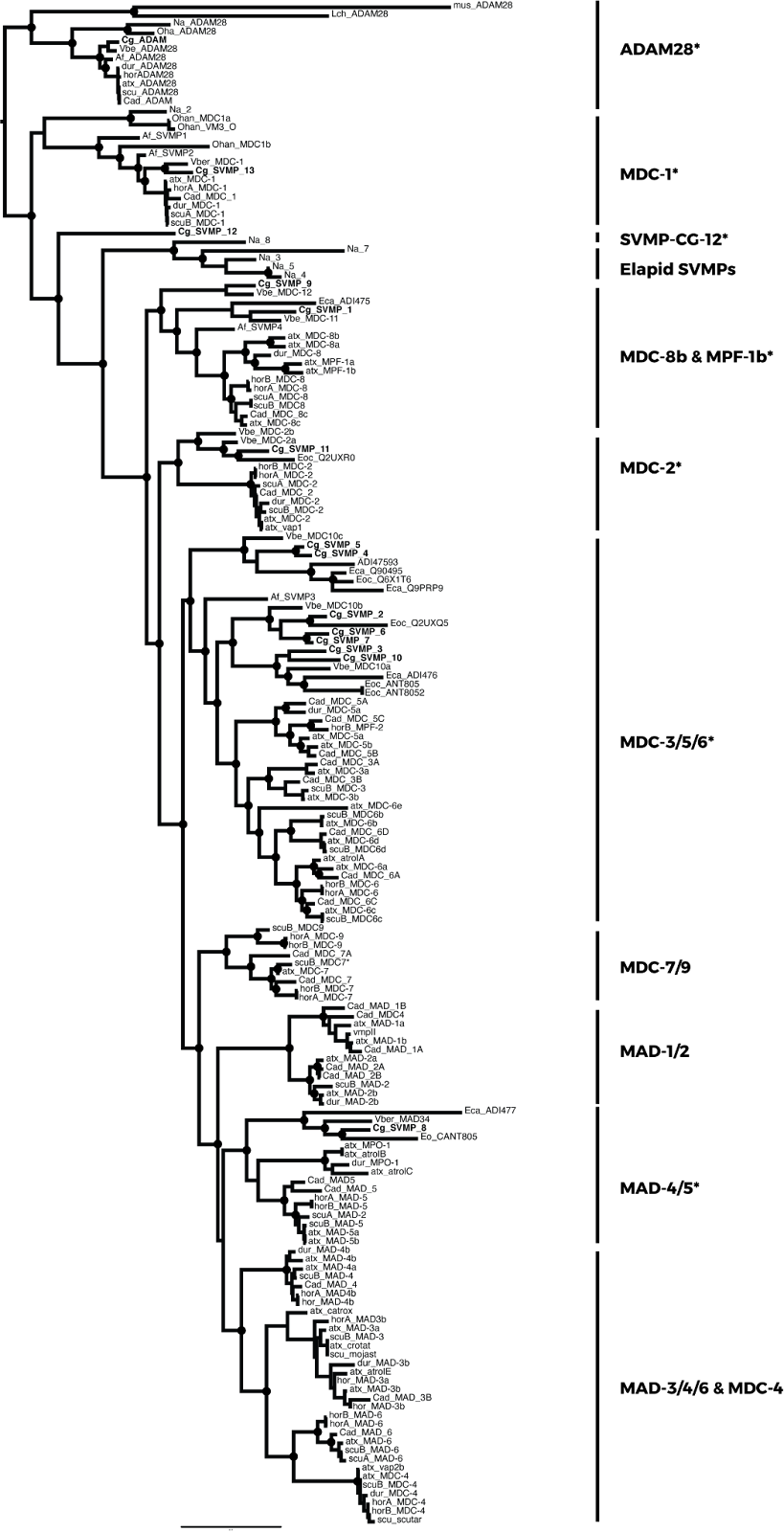

Fig. S6: Maximum likelihood phylogeny for SVMP genes and its non-toxic paralog (ADAM28). Genes for *Cerastes gasperettii* are highlighted in bold. Toxin groups are identified following previous categorizations. Asterisks indicate if *Cerastes gasperettii* genes are present in that specific group. Branch support with aBayes values higher than 90 are depicted as circles.

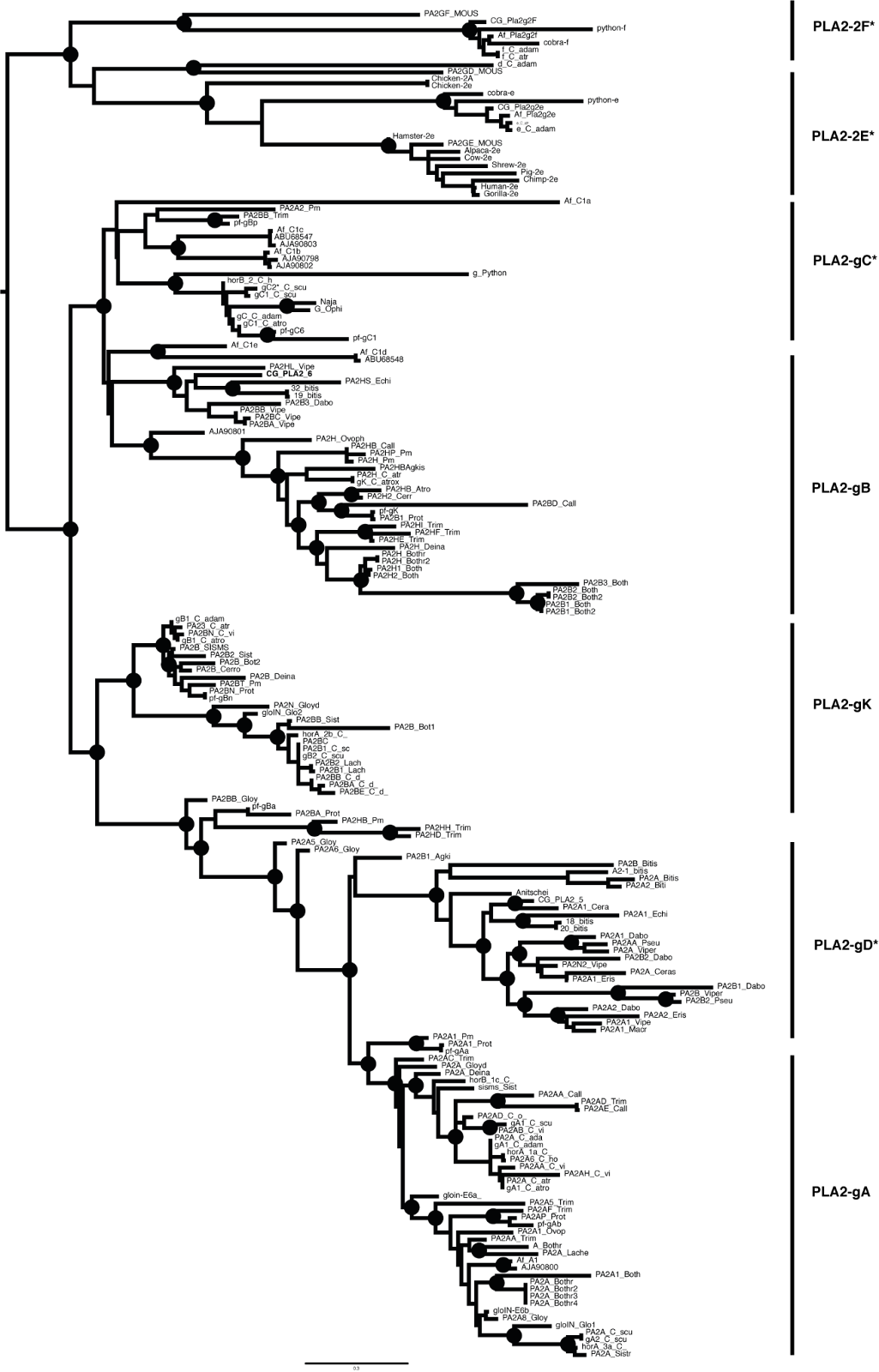

Fig. S7: Maximum likelihood phylogeny for PLA_2_, with the two non-toxic genes as outgroups (PLA_2_-2F and PLA_2_-2E). Asterisks in group labels indicate if *Cerastes gasperettii* genes are present in that specific group. Branch support with aBayes values higher than 90 are depicted as circles.

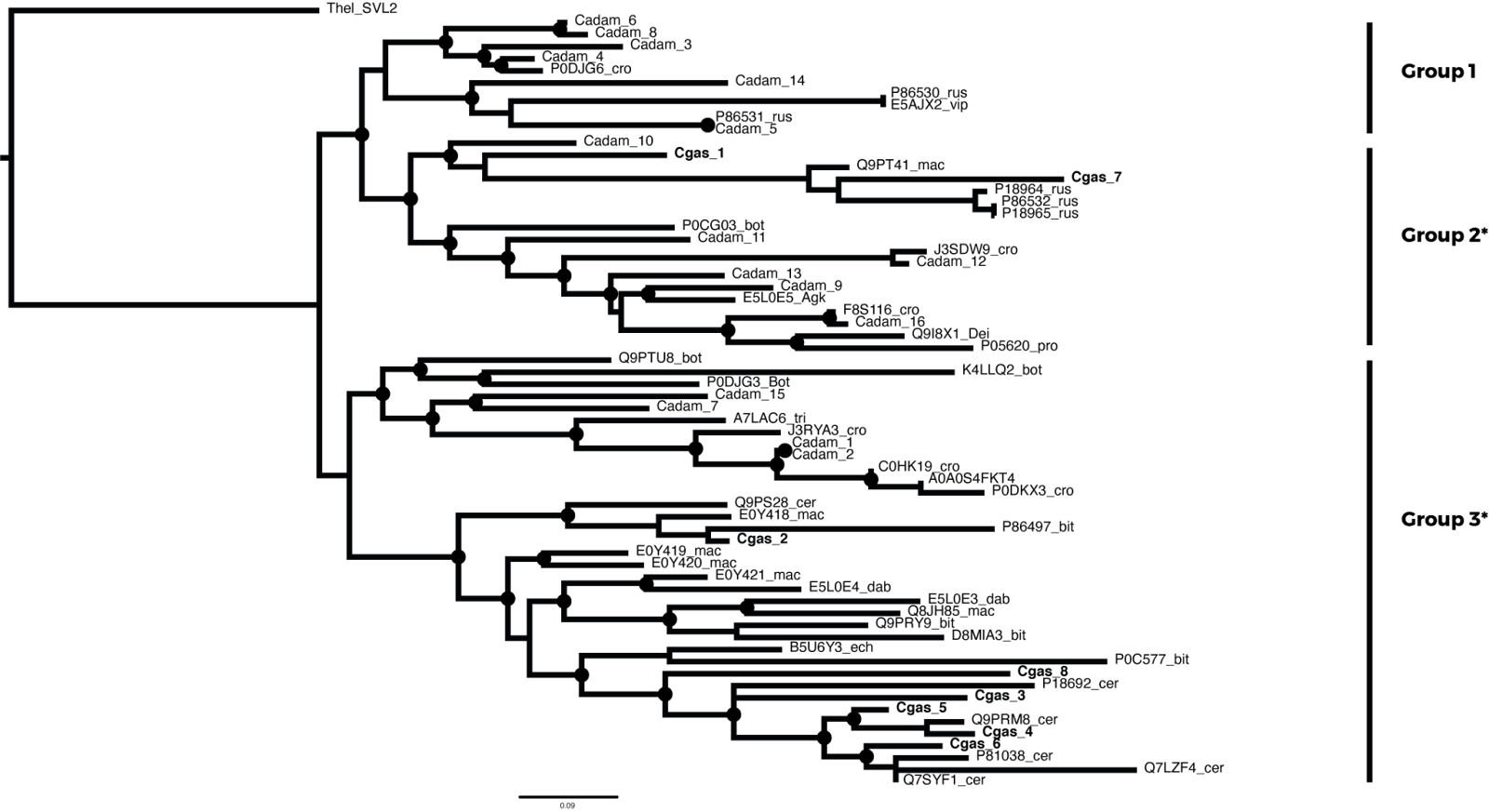

Fig. S8: Maximum likelihood phylogeny for SVSPs, with one sample from *Thamnophis elegans* as outgroup. Asterisks in group labels indicate if *Cerastes gasperettii* genes are present in that specific group. Branch support with aBayes values higher than 90 are depicted as circles.

Table S1: Individuals sampled in this study with their sex, sampling coordinates and data sequenced.

| **ID** | **Sex** | **Latitude** | **Longitude** | **Data sequenced** |
| --- | --- | --- | --- | --- |
| CG1 | Female | 25.284690 | 55.687860 | HiFi, Omni-C, Illumina, RNA-seq, Iso-seq, Proteome |
| CG9 | Female | 25.284690 | 55.687860 | RNA-seq |
| CG14 | Male | 25.284690 | 55.687860 | RNA-seq |
| CN6134 | - | - | - | Proteome |
| CN6135 | - | - | - | Proteome |

Table S2: Id, tissue type and number of reads sequenced per sample.

| **ID** | **Tissue** | **Reads** |
| --- | --- | --- |
| CG9 | Tongue | 44,672,733 |
| CG9 | Venom gland | 41,124,132 |
| CG9 | Eye | 41,951,109 |
| CG9 | Brain | 42,800,966 |
| CG9 | Heart | 40,715,947 |
| CG9 | Lung | 42,518,938 |
| CG9 | Liver | 42,251,137 |
| CG9 | Gallbladder | 42,738,665 |
| CG9 | Spleen | 40,909,550 |
| CG9 | Pancreas | 40,527,010 |
| CG9 | Ovary | 41,118,336 |
| CG9 | Kidney | 40,620,023 |
| CG9 | Accessory gland | 44,114,293 |
| CG14 | Tongue | 41,455,346 |
| CG14 | Venom gland | 41,035,764 |
| CG14 | Eye | 40,753,220 |
| CG14 | Brain | 43,413,973 |
| CG14 | Heart | 42,338,980 |
| CG14 | Lung | 42,068,410 |
| CG14 | Liver | 21,549,210 |
| CG14 | Gallglabbder | 50,571,941 |
| CG14 | Spleen | 45,447,235 |
| CG14 | Pancreas | 50,223,941 |
| CG14 | Testis | 47,495,900 |
| CG14 | Kidney | 45,945,776 |
| CG1 | Heart | 47,362,067 |
| CG1 | Brain | 45,740,571 |
| CG1 | Kidney | 50,758,869 |
| CG1 | Gallbladder | 40,546,711 |
| CG1 | Liver | 48,058,958 |
| CG1 | Spleen | 44,752,981 |
| CG1 | Tongue | 46,837,490 |
| CG1 | Pancreas | 45,023,783 |
| CG1 | Venom gland | 49,775,424 |
| CG1 | Ovary | 48,703,420 |

Table S3: Different types of repetitive elements masked within the genome:

| **Element** | **Number of elements** | **Length (bp)** | **Percentage** |
| --- | --- | --- | --- |
| Retroelements | 1524124 | 493932584 | 30.25 % |
| SINEs: | 339152 | 55265721 | 3.38 |
| Penelope | 124778 | 19471740 | 1.19 |
| LINEs: | 988815 | 347028895 | 21.25 |
| CRE/SLACS | 0 | 0 | 0.00% |
| L2/CR1/Rex | 480371 | 137654000 | 8.43 |
| R1/LOA/Jockey | 579 | 99034 | 0.01 |
| R2/R4/NeSL | 41793 | 10873028 | 0.67 |
| RTE/Bov-B | 128092 | 79663597 | 4.88 |
| L1/CIN4 | 207974 | 95913575 | 5.87 |
| LTR elements: | 196157 | 91637968 | 5.61 |
| BEL/Pao | 16545 | 5263265 | 0.32 |
| Ty1/Copia | 25582 | 15088781 | 0.92 |
| Gypsy/DIRS1 | 102598 | 63604234 | 3.90 |
| Retroviral | 50617 | 7642063 | 0.47 |
| DNA transposons | 707499 | 111444059 | 6.83 |
| hobo-Activator | 265944 | 30679712 | 1.88 |
| Tc1-IS630-Pogo | 227637 | 58877559 | 3.61 |
| En-Spm | 0 | 0 | 0.00% |
| MULE-MuDR | 44 | 3962 | 0.00% |
| PiggyBac | 138 | 6619 | 0.00% |
| Tourist/Harbinger | 182161 | 18395721 | 1.13 |
| Other | 0 | 0 | 0.00% |
| Rolling-circles | 2242 | 136656 | 0.01 |
| Unclassified | 205700 | 42385187 | 2.60 |
| Total interspersed repeats | - | 647761830 | 39.67 |
| Small RNA | 6134 | 652217 | 0.04 |
| Satellites | 35838 | 4217238 | 0.26 |
| Simple repeats | 765726 | 53044358 | 3.25 |
| Low complexity | 97863 | 6694649 | 0.41 |
